# Mesoscale Transport of Enveloped Viruses

**DOI:** 10.1101/2025.07.02.662892

**Authors:** Daniela Moreno-Chaparro, Florencio Balboa Usabiaga, Cecilia Zaza, David J. Williamson, Harry S. Holmes, Irene Carlon-Andres, Sabrina Simoncelli, Sergi Padilla-Parra, Marco Ellero, Nicolas Moreno

## Abstract

Enveloped viruses are characterized by spike proteins that protrude from and decorate the viral membrane. These proteins play a crucial role in host cell interactions and exhibit dynamic behaviors, such as tilting, sliding, and clustering, which vary across different types of enveloped viruses. For instance, SARS-CoV-2 spikes tilt to facilitate receptor binding, Influenza spikes migrate during infection, and HIV spikes migrate and cluster to enhance infectivity. In this study, we investigate how such dynamics influence the virus mobility. We characterize viral mobility through translational and rotational diffusion coefficients using a mesoscopic model that incorporates the dynamics of both the flexible spike proteins and the viral envelope. Using the smoothed dissipative particle dynamics (SDPD) method, we construct three virion models with varying spike flexibility. The first is a fully rigid virus with static spikes, the second is a model with spikes that tilt but remain fixed in position, and the third is a model allowing both tilting and sliding of spikes across the envelope. Our results show that spike flexibility primarily affects rotational diffusion, whereas the envelope dominates translational mobility of the virus. We also explore spike clustering driven purely by hydrodynamic interactions and compare with an experimental model reference using DNA-PAINT super-resolution imaging of HIV-like particles. We identify that hydrodynamic interactions alone can be responsible for dynamic clustering of spike proteins. Where, the characteristic size and lifespan of such clusters indicate predominantly doublets and triplets formations. Our findings highlight the role of spike dynamics in whole virion mobility, and motivate further investigations with time-resolved experimental evidence to fully characterize clustering behavior.

## I. INTRODUCTION

Viruses propagation and transmission have a significant impact on global-public health, as evidenced by the recent SARS-CoV-2 pandemic. Besides SARS-CoV-2, other enveloped viruses, such as HIV, herpes, dengue, and influenza have persisted for decades, continuing to cause fatalities and reduce quality of life worldwide. Enveloped viruses are characterized by a lipid bilayer membrane that encapsulates their genetic material. This membrane is decorated with spike proteins that play a crucial role in the virus’s ability to infect host cells. These spike proteins can bind to specific receptors on the cell surface, facilitating fusion and subsequent viral replication.^1^

Spike proteins can exhibit different dynamic behaviors that has been associated with the virus’s ability to infect host cells and evade the immune system. Depending on the virus type and the specific interactions involved, spike proteins can tilt, diffuse, and cluster over the viral envelope. For example, in SARS-CoV-2, the spike proteins have been reported to exhibit a tilting motion (ranging from 25^°^–30^°^) that enhance their ability to bind to host cell receptors.^2–4^ For Influenza viruses, the migration and localization of surface proteins has been identified as a hallmark during the infection process.^5–7^ In the case of HIV, besides tilting^8^ (ranging from 15^°^–25^°^) and diffusion of the surface proteins, recent studies suggested that the spike proteins can dynamically cluster, forming larger aggregates that can enhance the virus’s infectivity.^8–12^

Understanding the dynamic behavior of viral spike proteins is critical for interpreting the mechanisms of viral entry.However, at the scale of the entire virion, the influence of spike protein dynamics on viral transport remains largely unexplored. These conformational and fluctuating motions can modulate both translational and rotational diffusion, thereby impacting the passive mobility of the virus through the fluid medium. Viral diffusion is a multiscale process governed by a complex interplay of factors, including virion size and geometry, medium viscosity, and hydrodynamic interactions with the environment. Importantly, thermally driven fluctuations in the spike proteins can alter mesoscale transport properties of the virion.

The coupling between microscopic spike protein dynamics and mesoscale virion mobility presents a fundamental challenge from both experimental and computational perspectives. Experimentally, a variety of advanced techniques have been employed to probe the structural, mechanical, and functional properties of enveloped viruses.^9,13–19^ While these tools have yielded valuable insights into virus assembly, deformation, and spike-mediated host interactions, they fall short of capturing the dynamic interplay between spike conformational changes and whole-virion transport. The simultaneous resolution of molecular fluctuations and mesoscale motion in fluid environments remains a formidable challenge.^20^ On the computational front, numerical modeling has become indispensable in bridging these gaps.^21,22^ Molecular dynamics (MD) simulations^23–26^ offer atomistic-level resolution of viral structures and interactions, yet they are often limited by system size and timescale constraints. To overcome these limitations, a range of mesoscale modeling approaches has been developed, including Brownian dynamics,^27,28^ coarse-grained (CG) models,^29,30^ dissipative particle dynamics (DPD),^31–33^ Monte Carlo simulations,^34^ rigid bead-rod theory,^35,36^ and rigid multi-blob (RMB) models.^37–39^ While these methods enable the exploration of larger spatiotemporal scales and capture hydrodynamic effects, integrating fluctuating spike dynamics with realistic virion geometry and fluid-mediated interactions remains to be fully realized.

Recently, Moreno et al^38^ adopted a rigid multi-blob model to characterize the translational and rotational diffusion of different enveloped viruses, exploring the effects of spike-proteins density and morphology. The viruses were modelled as fully-rigid objects, and their passive transport was associate solely on geometrical features. The authors postulated the existence of an optimal balance between viral rotational diffusion and interaction, modulated by the number of spike proteins.^38^ The optimal condition showed to hold for the majority viruses investigated. However, viruses such as HIV, did not comply with this trend. This discrepancy suggested that the rigid models may not fully capture the effect of complex dynamics of HIV spike proteins^9–12^ on the overall virion transport. This limitation of fully-rigid models motivates the development of more realistic virion models, where the translational degrees of freedom of the spike-proteins are relaxed.

Herein, we aim to develop a generalized mesoscopic model of spike/virion diffusion that bridges experimental and theoretical insights across spatial and temporal scales. To this end, we construct mesoscale models of enveloped virions suitable to study their diffusion, while accounting for spike-proteins dynamics. We model the hydrodynamic interactions between the virion and the surrounding fluid using the Smoothed Dissipative Particle Dynamics^40,41^ (SDPD) method. We extend the rigid represention of the virions used by Moreno and coauthors,^38^ by modelling the spike proteins and envelope as separate rigid bodies. This allow us to account for various spike proteins dynamics such as tilting and diffusion over the envelope. We analyze the change in the translational and rotational diffusion of the virions, with the variations in the spike proteins dynamics. Additionally, given the relevance of spike proteins clustering in viral infectivity,^9–12^ we explore the applicability of our mesoscale virion models to capture clustering behavior of spike proteins, and use super-resolution imaging of HIV-1 particles decorated with JR-FL v4 ALFA-tag with the C terminus truncated (Delta CT); which means that spike-proteins diffuse freely over the viral surface without being anchored to the underlying matrix, and their incorporation into single viruses is not regulated. We qualitatively compare the clustering behavior for our mesoscale virion models and the images of HIV-1 particles and assess whether similar dynamics emerge in our simulations. This provides a relevant experimental context for evaluating our model predictions to compare clustering dynamics when proteins can freely diffuse over the envelope, mainly dominated by hydrodynamic interactions.

In the remainder of this document, in Section II, we present the discretization and modeling of the virions. In Section III, we present the diffusional results for virions with different types of spike proteins dynamics, and discuss the implications of our finding in the transport properties of enveloped viruses. We finally explore the potential formation of spike proteins clusters mediated by hydrodynamic interactions, and compare with experimental evidence observed in virion particles.

## II. METHODS AND MODELS

### A. Smoothed Dissipative Particle Dynamics

SDPD is a mesoscale particle-based method widely used to model complex fluids, incorporating consistent thermal fluctuations of the dissipative particle dynamics^40,41^ method and the discretization scheme of smoothed particle hydrodynamics.^42^ SDPD has been successfully applied to model a wide range of complex systems, including synthetic^38,43^ and biological^44,45^ polymeric materials, multiphase flows,^46^ colloids,^47^ and red blood cells to name a few. In this framework, we discretize both the virus and the surrounding fluid into small spherical particles. The SDPD method incorporates both conservative and dissipative forces, as well as random thermal fluctuations, to model the interactions between particles. Each particle is characterized by its position **r**_*i*_, velocity **v**_*i*_, mass *m*_*i*_, and a volume *V*_*i*_, such that 1*/V*_*i*_ = *d*_*i*_ = ∑ _*j*_ *W* (*r*_*i* *j*_, *h*). Here *d*_*i*_ is the number density of particles, *r*_*i* *j*_ =|**r**_*i*_−**r** _*j*_|, and *W* (*r*_*i* *j*_, *h*) is an interpolant kernel with finite support *h*. In 3D, we use the kernel *W* (*r*) = 105*/*(16*πh*)(1 + 3*r/h*)(1 − *r/h*)^3^ if *r/h <* 1 or *W* (*r*) = 0 if *r/h* ≥ 1.^40^

The evolution equations for the particles position is d**r**_*i*_/*dt* = **v**_*i*_, whereas the momentum equation takes the form

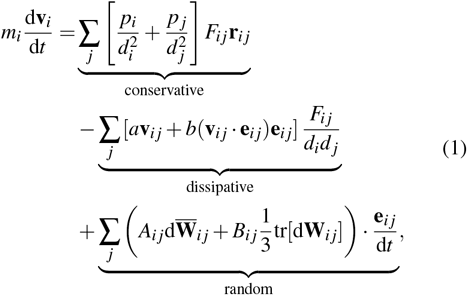

where **e**_*i* *j*_ = **r**_*i* *j*_*/*|**r**_*i* *j*_|, **v**_*i* *j*_ = **v**_*i*_−**v** _*j*_, *a* and *b* are friction coefficients related to the shear *η* and bulk *ζ* viscosities of the fluid through *a* = 5*η/*3 − *ζ* and *b* = 5(*ζ* + *η/*3). In Eq. 1, we introduce the positive function *F*_*i* *j*_ = −∇*W* (*r*_*i* *j*_, *h*)*/r*_*i* *j*_. The term 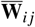 is a matrix of independent increments of a Wiener process for each pair *i, j* of particles, and 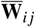 is its traceless symmetric part, given by 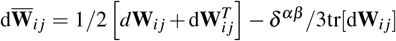 To satisfy the fluctuation-dissipation balance the amplitude of the thermal noises *A*_*i* *j*_ and *B*_*i* *j*_ are related to the friction coefficients *a* and *b* through *A*_*i* *j*_ = [4*k*_*B*_*TaF*_*i* *j*_*/ρ*_*i*_*ρ*_*j*_]^1*/*2^ and *B*_*i* *j*_ = [4*k*_*B*_*T* (*b*−*a/*3)*F*_*i* *j*_*/ρ*_*i*_*ρ*_*j*_]^1*/*2^, where *k*_*B*_*T* is the thermal energy, and *ρ*_*i*_ and *ρ*_*j*_ are the densities of the particles *i* and *j*.The pressure *p*_*i*_ of the *i*-th particle is estimated using an equation of state of the form *p*_*i*_ = *c*^2^*ρ*_0_*/*7[(*ρ*_*i*_*/ρ*_0_)^7^ −1], where *c* is the artificial speed of sound on the fluid, and *ρ*_0_ is the reference density. The term *c*^2^*ρ*_0_*/*7 sets the speed of sound at the reference pressure, *c*^2^ = *∂ p/∂ρ*|_*ρ*=ρ0_.

### B. Mesoscale Virion Models

We construct three mesoscale virion models to investigate the effect of spike proteins dynamics over the passive transport of enveloped viruses. These models corresponds to a spherical envelope decorated with (1) rigidly connected spikes proteins,(2) fixed-position pivoting spikes proteins, and (3) freely-sliding/pivoting spikes proteins, as illustrated in FIG. 1a.

**FIG. 1.**
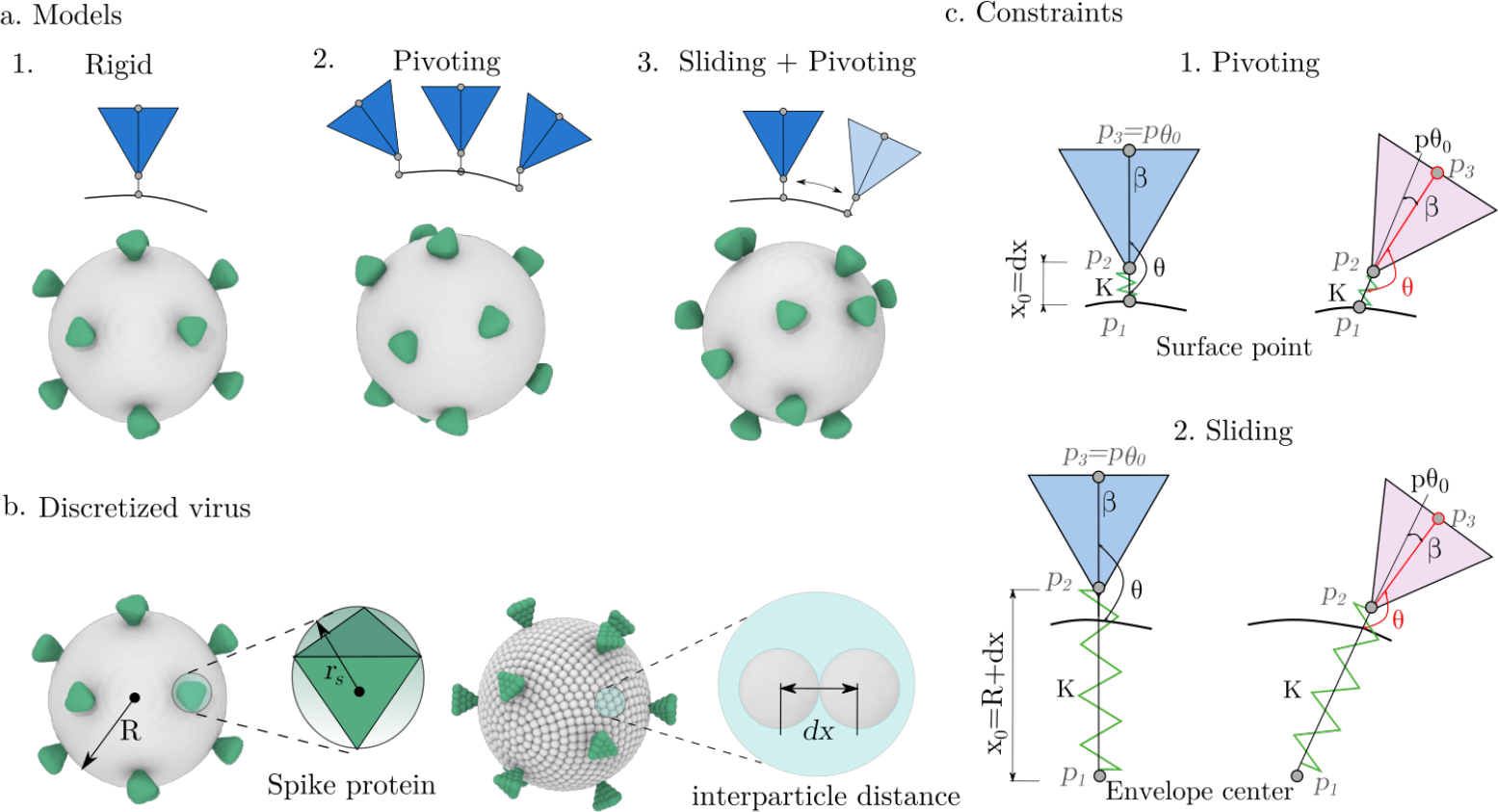
(a.)Schematic representation of the three virus models. spikes proteins are modeled as tetrahedral structures and are uniformly distributed over a spherical envelope. Model 1 features rigid spikes proteins, model 2 allows pivoting (tilting) motion, and model 3 incorporates both pivoting and sliding (free diffusion). (b) Discretization of a virion with envelope radius *R* and tetrahedral spike-protein radius *r*_*s*_, in SDPD. *dx* denotes the interparticle distance. (c.) Linear harmonic potential constraint, here, spring constant *K*, anchors each spike-protein base to the envelope. The distance *x*_*se*_ is measured between the points | **p**_1_ −**p**_2_ |. The spring equilibrium distance is *x*_0_ = *dx* for the pivoting case (c.1)and *x*_0_ = *R* + *dx* for the sliding case (c.2). For the angular harmonic potential constraint, the spring constant *K*_*α*_ controls the stiffness in three points **p**_1_, **p**_2_, **p**_3_. Here, we define *θ* as the angle formed between these three points, with an equilibrium angle *θ*_0_ = 180. Blue spike protein illustrate *θ* = *θ*_0_, when **p**_3_ = **p***θ*_0_. Tilting angle *β >* 0 amounts for deviations from the equilibrium case (pink spike proteins).

#### 1. Model 1

In Model 1, the whole virion (envelope and spike proteins) is represented as a rigid body, where the spike proteins are uniformly distributed over the surface of the envelope. This initial approximation serves as a reference point for comparing the effects of spike flexibility in subsequent models, and allows us to compare with the prior diffusional studies by Moreno et al,^38^ where the surrounding fluid was modeled implicitly using the Rigid Multi-Blob (RMB) method.

#### 2. Model 2

In Model 2, we assess how spike tilting affects the whole-virion diffusion. Here, spike proteins are represented as independent rigid bodies and are allowed to pivot at fixed positions over the envelope. Two additional potentials govern spike-envelope interactions. First, the spike protein is anchored to a fixed position over the envelope surface (as occur in model 1), *via* an harmonic spring potential, *E* = (1*/*2)*K*(*x*_*se*_−*x*_0_)^2^, where *K* is the spring constant, *x*_*se*_ =|**p**_1_−**p**_2_|the distance between two prescribed particles in the spike protein and the envelope respectively, and *x*_0_ = *dx* is the equilibrium distance of the potential, defined to be near the equilibrium SDPD interparticle distance (*dx*), to prevent artificial density variations.

MD studies often describe spike proteins tilting using multi-joint structure^2^ (hip-knee-foot) with cumulative articulation angles. Here, we adopt a single pivot located at the base of each spike proteins describing an effective tilting angle with respect to the envelope’s surface, *via* an harmonic angular spring potential. This potential is given by *E*_*α*_ = (1*/*2)*K*_*α*_ (*θ*−*θ*_0_)^2^, where *K*_*α*_ is a constant that controls the stiffness, *θ*_0_ is the equilibrium angle, and *θ* is the angle formed between a fixed point in the envelope surface (**p**_1_),a point in the protein base (**p**_2_), and the protein tip (**p**_3_), as illustrated in Fig. 1.c.1.

In our virions, we define the tilting angle *β* =|*θ*−*θ*_0_|of the spike proteins to prescribe a favored orientation, and *K*_*α*_ tunes the degree of flexibility of the protein with respect to this orientation. Here, we set an equilibrium angle *θ*_0_ = 180^*°*^, corresponding to a perpendicular alignment over the envelope surface. In the limit *K*_*α*_→∞, the spike protein remains perpendicular to the surface fixed point, *β*→0. Whereas, as *K*_*α*_ *<* ∞, we expect *β >* 0 as the thermal fluctuations become relevant. In the results section, we show that the magnitude of *β* can be effectively adjusted by varying *K*_*α*_. Additionally, we compare with the theoretical expectation derived from the probability distribution of the intrinsic tilting angle *β* (See SI, III, Table II). Given the angular potential adopted, the distribution function reads *P*(*β*) = *Z*^−1^ sin(*β*) exp(-*K*_*α*_ (*β*)^2^*/*2*k*_*B*_*T)*, where *Z* is a normalization constant that ensures the integral of the probability function is equal to one, 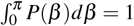 1. This expression enables estimation of the expected values of *β* as a function of *K*_*α*_, providing a theoretical baseline for comparison with our simulation results.

#### 3. Model 3

In Model 3, the spike proteins can both pivot and slide over the envelope surface. This configuration introduces additional degrees of freedom associated with the free diffusion of spike proteins over the envelope’s surface. Similar to Model 2, we use two harmonic potentials to constraint the position and tilting. In this case, the spike proteins are anchored to the envelope center of mass, **p**_1_, with an harmonic potential at an equilibrium distance *x*_0_ = *R* + *dx*, where *R* is the envelope radius, as illustrated in FIG. 1.c.2. The control over the tilting angle is modeled using an harmonic angular potential, where the angle *θ* is formed between the envelope’s center of mass **p**_1_, a point in the protein base (**p**_2_), and the protein tip (**p**_3_), as shown in FIG. 1.c.2. The equilibrium angle is *θ*_0_ = 180 ^*°*^, ensuring an alignment of the spike protein normal to the envelope’s center of mass. For model 3, the hydrodynamic interactions with the surrounding fluid can induce both translation and tilting of the spike proteins. Since we are not explicitly accounting for translational constraints (besides fluid drag), we vary the surface protein stiffness to investigate scenarios where either sliding or tilting dynamics can be dominant.

Moreover, we explore a variant of Model 3 where the spike proteins move over a fixed envelope to simulate membrane-bound behavior. Thus, we also compute the geodesic mean-squared displacement, *gMSD* of the spike proteins (see SI, VIII for detailed description) to estimate their effective translational diffusion *D*_*sp*_ over the envelope. In principle, since *D*_*sp*_ can be affected by the amount of decorating proteins, we vary the numbers *N*_*sp*_ of spike proteins, ranging from 2-24 per envelope, to investigate the presence of a crowding effect.

### C. Whole-Virion Passive Transport

To analyze the impact of spike-protein dynamics on virus mobility, we compute the translational (*D*_*t*_) and rotational (*D*_*r*_) diffusion coefficients using the mean square displacement, MSD = 6*D*_*t*_*t* (see SI, VII) and mean square angular displacement, MSAD = 4*D*_*r*_*t* (see SI, VI), respectively.^47–49^ For the each of the models evaluated, we perform three independent realizations to ensure statistical robustness. *D*_*t*_ and *D*_*r*_ are estimated from the MSD in the range 10*τ*_*ν*_ to 100*τ*_*ν*_, where 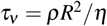 is the kinematic time, *ρ* is the density mass, *R* is the envelope radius and *η* the viscosity^50^.

While factors such as size, shape, number, and distribution of spike proteins influence virion transport,^38^ this study focuses solely on the effects of spike dynamics. For generality, we model enveloped virions with a spike-to-envelope size ratio of *r*_*s*_/*R* ≈ 1*/*10, where *r*_*s*_ is the spike radius. Inspired by HIV and SARS-CoV-2 structures, we represent spikes proteins as tetrahedral-like shapes, with *r*_*s*_ defined as the radius of the sphere inscribing the tetrahedron. Unless otherwise stated, we use a fixed number of twelve uniformly distributed spikes proteins (see FIG. 1.b), consistent with experimental observations of HIV virions.^38^

### D. Spike-proteins Clustering

We investigate the occurrence of dynamic spike-proteins clustering mediated by hydrodynamic interactions in model type 3, where the spike proteins can freely diffuse over the envelope. When any particle belonging to one spike protein is within a clustering cutoff radius *r*_*cl*_ = 5*r*_*s*_, of a particle from another spike protein, we consider the spike proteins form part of the same cluster. To monitor the temporal evolution of cluster formation throughout the simulation, we perform a clustering analysis on simulation snapshots taken every 1000Δ*t*. The total number *N* of clusters and number *C*_*s*_ of spike proteins per cluster is measured to estimate the *lifetime l*_*t*_(*C*_*s*_). The lifetime *l*_*t*_(*C*_*S*_) is the cumulative time (not necessarily consecutively) that clusters of size *i* are observed during the simulations. We rationalize the relevance of the times scales of spike-proteins clustering events, with respect to the characteristic diffusive time *τ*_sat_ = *R*^2^*/D*_*sp*_ of the spike proteins over the envelope surface. Thus we define the normalized lifetime

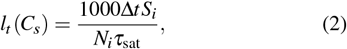

where *S*_*i*_ is the total number of snapshots that a cluster with size *C*_*s*_ = *i* appear in the simulation, and *N*_*i*_ is the number of such clusters. We also define the *lifespan* Δ*s*(*i*) as the number of consecutive snapshots during which cluster of size *C*_*s*_ remains. Further reappearances are treated as new lifespan event. Thus, the mean lifespan is given by

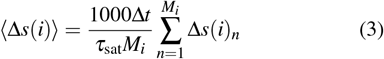

where *M*_*i*_ is the total number of lifespan events observed during the simulation for a cluster of size *C*_*s*_ = *i*.

### E. Simulation Setup and Parameters

As is customary in SDPD simulations, we adopt a system of dimensionless (reduced) units by defining characteristic scales for length (*l*_sdpd_), mass (*m*_sdpd_), and energy (*ε*_sdpd_). Consequently the characteristic time unit is *τ*_sdpd_ = *l*_sdpd_(*m*_sdpd_*/ε*_sdpd_)^1*/*2^. All quantities are expressed in terms of these fundamental units. In all the simulations,we use periodic cubic domains with size *L*_*b*_ = 15*l*_sdpd_. The interparticle distance is set to *dx* = 0.2[*l*_sdpd_], leading to an equilibrium particle density 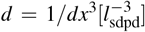, whereas the size of the interpolation kernel is *h* = 4*dx*, ensuring that hydrodynamic interactions are accurately captured while maintaining computational efficiency (see SI, II).^42,50,51^

In our discretization, the envelope and spike proteins are formed by SDPD particles located with an interparticle distance *dx* = (1*/d*)^1*/*3^, where *d* is the particle number density of the simulation. We define the resolution of the virion model from the ratio *R/dx*, where *R* is the radius of the envelope. To determine an appropriate resolution, we conduct a convergence study (see SI, II) examining the rotational mobility of spherical envelopes. We find that a resolution of *R/dx* = 10 offers an optimal trade-off between computational efficiency and consistency with theoretical predictions from Stokes-Einstein theory.^52^ This choice ensures that the envelope is sufficiently resolved to capture both its translational and rotational dynamics while maintaining tractable computational cost for simulations involving the spike proteins. At this resolution, we quantify the numerical deviations of the simulated envelope diffusion coefficients (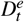 and 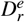) relative to theoretical Stokes–Einstein values: approximately 6% for translational and 24% for rotational diffusion. The larger deviation in the rotational component is attributed to its *R*^3^ dependence, which enhances sensitivity to finite-size effects. However, considering that these numerical deviations are systematic and persist across all virion models considered, we follow the approach of Moreno et al,^38^ introducing reduced diffusion coefficients

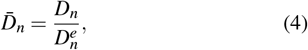

where *D*_*n*_ is the measured diffusion coefficient of a virion model (including spikes proteins), 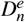 is the corresponding value measured for the bare spherical envelope (without spikes proteins), and *n* = *t, r* denotes the translational or rotational component, respectively. Eq.(4) allows us to normalize out numerical deviations and enable direct comparison between models. In the results section, we show that for the virion resolution adopted, we reproduce the rotational diffusion 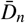 previously reported for fully rigid models.^38^

For virion models with mobile spike proteins, we set the spring constant 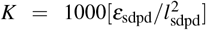, whereas different angular spring constant are explored *K*_*α*_ = {8, 10, 100, 1000}[*ε*_sdpd_].

### F. Spike-protein clustering in virion-like particles

We support our computational analysis by conducting DNA-PAINT^53^ super-resolution imaging of mutated HIV-1 virion particles^54^ and examining the organization and clustering of spike proteins on the viral surface (see SI, IX for details). The virion particles are decorated with HIV-1 spike proteins (a.k.a *Env proteins*) and are comparable in size to native HIV-1 virions. However, the spike-proteins exhibit a mutation^54^ in the variable 4 (v4) loop to allow single domain nanobodies to engage with the ALFA-tag docking site. Also, the C terminus of the spike protein was truncated to allow them to freely diffuse over the virus surface, making this virus a valuable experimental reference for interpreting the clustering behavior explored in Model 3. We must note that, since the virion particles need to be immobilized, the temporal evolution of the clusters cannot be directly estimated. However, this technique provides a 2D-staggered projection of the virions, allowing us to assess quantitative information about cluster size and distribution.

We immobilize fixed virion particles on a functionalized glass surface and use nanobodies labeled with short singlestranded DNA to target specific surface proteins. These DNA strands enabled high-precision imaging through sequential binding of fluorescently labeled complementary strands in solution. Then, using a custom-built TIRF microscope (SI, IX.F), we acquire tens of thousands of images and reconstruct super-resolved images by localizing single-molecule events. Gold nanoparticles serve as fiducial markers for drift correction and spatial accuracy. Finally, we use a quantitative analysis method (qPAINT)^53,55^ to obtain 2D estimates of the number of proteins per cluster and identify patterns (SI, IX.H).

## III. RESULTS AND DISCUSSION

### A. Model 1: Fully-Rigid Model Diffusion

In FIG. 2a, we compile the MSD and MSAD curves (averaged over three realizations) for fully-rigid models. The linear variation of MSD consistently indicates a normal diffusion behavior of the virion. Similarly, MSAD exhibits normal diffusion at short times, but eventually saturates near the

**FIG. 2.**
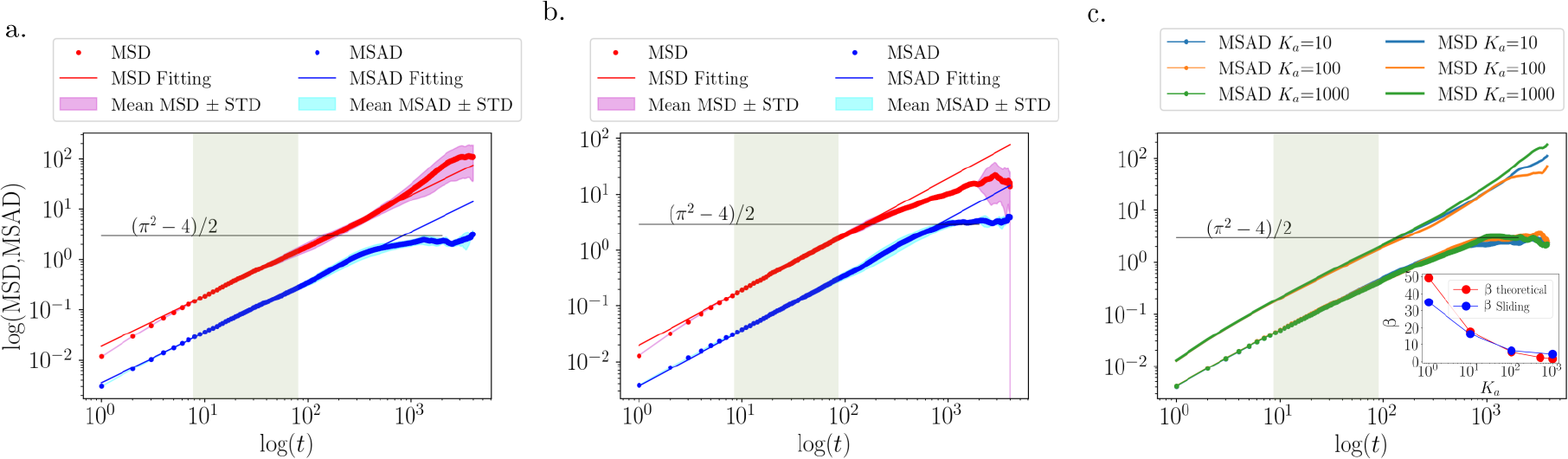
a. MSD and MSAD log-log curves for the rigid virion models averaged over three independent realizations. The shaded green region marks the interval used to compute rotational diffusion 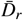, and translational diffusion 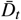. The MSAD curve saturates near the theoretical limit (*π*^2^ −4)*/*2. b. MSD and MSAD log-log curves for pivoting model and c. MSD and MSAD log-log curves for sliding models respectively for three different cases of *K*_*α*_. We use the continuous lines for the MSD and the continues with dots for the MSAD. Inside plot, show the tilting angle *β* as a function of angular stiffness *K*_*α*_ (1 to 1000) for both theoretical predictions and simulation results on curved (envelope) surfaces. *β* is defined between three points: envelope center of mass (*p*_1_), spike base (*p*_2_), and spike tip (*p*_3_). Higher *K*_*α*_ values restrict spike motion, reducing *β*. For the most flexible case, *K*_*α*_ = 1, the theoretical and simulated angles differ because, in the simulation, the tetrahedral spike can collide with the envelope, diminishing the effective angle inclination.

expected theoretical^49^ bound (*π*^2^ −4)*/*2 due to angular confinement. In FIG. 2, the shaded green (10*τ*_*ν*_ to 100*τ*_*ν*_) region highlight the intervals used to compute 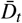 and 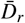. The normalized diffusion coefficients obtained, 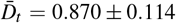 and 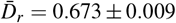, evidence a reduction in both translational and rotational diffusivity compared to the bare envelope. Remarkably, this results are in close agreement with the measured diffusions using RMB^39^ with 12 uniformly distributed spike proteins (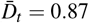 and 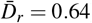). These results support the accuracy of our SDPD implementation and the chosen resolution of the virion models.

### B. Model 2: Pivoting Model Diffusion

We now examine the diffusion behavior of the virion models featuring flexible pivoting spike proteins. In FIG. 2.b, we present the MSD and MSAD curves averaged over three independent simulations for virions with tilting dynamics, with a mean tilting angle *β*≈20^°^. Both curves exhibit a similar trend as the rigid model, with MSD showing linear growth and MSAD saturating near the theoretical limit (*π*^2^− 4)*/*2. The computed normalized diffusion coefficients for this model are 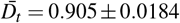 and 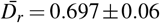. Compared to the rigid virion model, the pivoting configuration has a slightly larger translational and rotational diffusion. This indicates that the tilting dynamics of the spike proteins do not hinder significantly virus mobility.

### C. Model 3: Sliding and Pivoting

In FIG. 2.c, we compile the MSD and MSAD curves for the sliding model, varying the stiffness constant on the range *K*_*α*_ = {10, 100, 1000}[*ε*_sdpd_] (see also SI, IV Table III)for the summarized values of 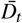 and 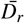). In the inside plot of FIG. 2.c, we also show the dependency of the measured tilting angle *β* with *K*_*α*_, to illustrate versatility of the model to reproduce a range of tilting angles. Consistent with theoretical predictions, increasing *K*_*α*_ reduces *β*, with angles approaching 0^*°*^ at the highest stiffness evaluated (*K*_*α*_ = 1000*ε*_sdpd_). Conversely, at the lowest *K*_*α*_ = 1*ε*_sdpd_, the tilting angle can reach a mean angle 50^*°*^, indicating nearly unconstrained tilting.

In FIG. 2.c, we can observe that the translational diffusion remains relatively stable across different *K*_*α*_ values, with 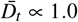, suggesting modest alterations with respect to the bare envelope, with variations falling within the range of simulation uncertainty. The slightly higher translational diffusion compared to the previous models, suggests that the spike proteins mobility may induce marginal gains in the translational motion, due to dynamic redistribution of hydrodynamic stresses. Regarding the rotational diffusion, virions with more flexible spike proteins (lower *K*_*α*_) exhibit a slightly larger rotational mobility. This is consistent with previous observations in pivoting models, where the flexibility appears to reduce rotational drag.

### D. Summary of Different Models and Implications

In FIG. 3, we summarize the variation of the translational and rotational diffusion for the free envelope and the different virion models investigated. We can identify that the dynamics of spike proteins over the envelope, are responsible for variations in 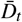, characterized by reductions around 10% for models with immobile spike proteins (model 1 and 2, rigid and pivoting, respectively). In contrast, virion model 3, with freely sliding spike proteins, exhibit a 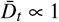 on the order of the free envelope. Moreover, FIG. 3.a shows that the tilting degree of the proteins induce only marginal variations in the translational diffusion of the virions, as evidence on the weak dependence of 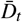 with *K*_*α*_.

**FIG. 3.**
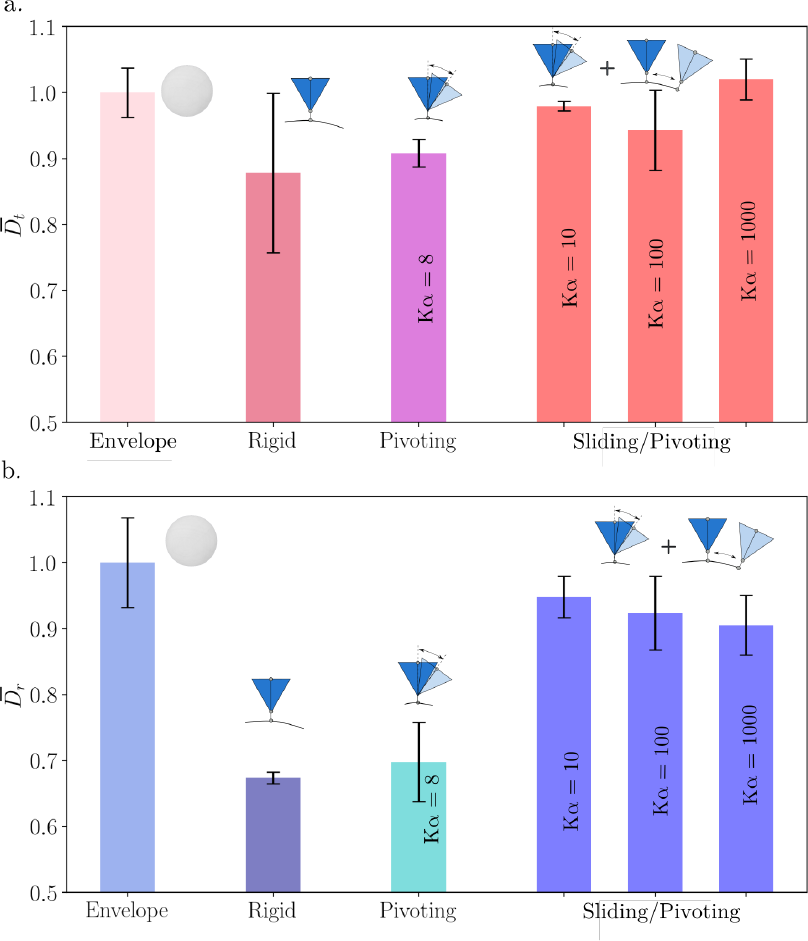
Normalized translational 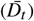 (a.) and rotational 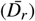 (b.) diffusion coefficients for the three virus models: rigid, pivoting, and sliding. For the rigid model, *K*_*α*_ →∞; for the pivoting model, *K*_*α*_≈8; and for the sliding model, *K*_*α*_ takes values of 10, 100, and 1000.First bar in both graphics indicate the baseline envelope diffusioncoefficients for translation and rotation, respectively.

In general, the impact of spike protein flexibility is small for the translational diffusion. However, for rotational diffusion spike proteins dynamics becomes more relevant. In FIG. 3.b, we can observe that both tilting and diffusion of the spike proteins alter the overall rotational motion of the virions. When comparing Models 1 (fully rigid) and 2 (immobile but freely tilting proteins), where spike proteins have a fixed position over envelope, the spike-protein tilting dynamics only induce a marginal increase in the rotational diffusion. In contrast, Model 3, where spike proteins can slide, exhibits the largest 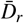. This suggest that the sliding of the spike proteins can facilitate the dynamic redistribution of hydrodynamic stresses leading to an overall reduction in the rotational drag. Additionally, as the spike protein becomes more flexible (i.e. decreasing *K*_*α*_), the marginal gains due to tilting dynamics, lead to 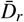 values converging to the bare envelope, indicating that envelope and spike-proteins mobility are decoupled.

Stemming from our current findings, we explore now the implications of 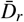 variations on the *excess diffusion* proposed by Moreno and coauthors.^38^ The authors defined an excess rotational diffusion Δ*D*_*r*_, given by 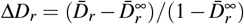, where 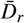 is the measured virion’s rotational diffusion with a fixed number of spike proteins, and 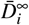 is the asymptotic value in diffusion coefficient, as the number of spike proteins reaches 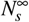, that is the saturation number of spikes. Thus, further addition of spike proteins over the envelope does not affect their mobility. Moreno and coauthors, postulate that for a given virion (e.g. HIV, SARS-CoV-2, Influenza), the variations in the excess diffusion can be expressed in terms of the total number of spike proteins, Δ*D*_*i*_ = *f* (*N*_*s*_), as

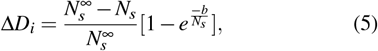

where 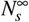 is in general defined by the aspect ratio between the spike protein and the envelope, and *b* indicates the rate of decay in the rotational mobility. In Eq.(5), *b* is a fitting parameter characteristic for each type virus, and lower values indicate a faster decay of Δ*D*_*i*_.

Models 2 and 3 can be associated with HIV virions with different state of maturation, where Model 2 corresponds to immature states, where the spike proteins are linked to the membrane and cannot diffuse^10,54,56,57^. Whereas Model 3 are consistent with mature states, when the capsid of the virus is already form and the spike proteins can translate along the membrane. We evaluate Eq. (5) for the three virion models,using a saturation value of 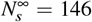 consistent for HIV.^38^ We estimate the decay parameters to be *b*_1_ = 5.8, *b*_2_ = 6.7, and *b*_3_ = 25.7, where the subindex denotes the model type. In FIG 4, we present Δ*D*_*r*_ as a function of *N*_*s*_, and indicate the corresponding excess diffusion at *N*_*s*_ = 12. This number falls within the experimentally reported range for HIV, typically between 11 and 14. At such low spike densities, we observe that the transition from a mature to an immature state can significantly impact the rotational motion of the virion. This raises the question of whether the maturation state itself could serve as a regulatory feature to modulate transport properties. It is important to note that friction between the spike proteins and the membrane can influence their mobility, thereby affecting the overall diffusion of the virion. In our framework, Model 2 and Model 3 represent the limiting cases of spike diffusion—fully mobile and fully immobile spikes, respectively—with real systems likely exhibiting behavior somewhere between these extremes. For virions with a larger number of spike proteins, such as herpes virus, or for larger virions in general, the differences between these two models may become negligible, as the influence of individual spike mobility on overall transport diminishes.

**FIG. 4.**
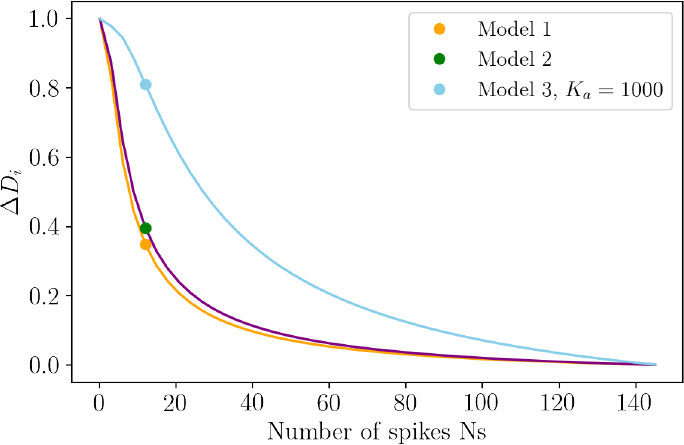
Comparison rotational diffusion (Δ*D*_*r*_) as a function of spike number (*N*_*s*_) for flexible and rigid virus model. The solid orange curve represents the rigid virus model, while the purple and blue curves shows our flexible models. Here for model 2 and 3, *b* values are 6.7 and 25.7 respectively. Markers on all curves indicate values for 12 spikes proteins. A clear increase the HIV sliding model 3, highlighting the impact of spike flexibility on rotational diffusion.

### E. Spike-Proteins Clustering Dynamics

Besides the effects of spike-protein dynamics on the whole-virion transport, we now use the mesoscale virion models to investigate the clustering dynamics of the spike proteins and compare with the 2D-staggered projections of HIV-like particles using DNA-PAINT super-resolution imaging.

In FIG. 5, we compare the gMSD of the spike proteins over envelopes containing *N*_*s*_ = 2, 12, 24 proteins. In FIG. 5.a, we observe that the gMSD curves exhibit linear dependency at short times and saturate at long times *t > t*_sat_ = *R*^2^*/D*_*sp*_. The simulated virions show an equivalent behavior at short times and consistently saturate to values close to *R*^2^(*π*− 4)*/*2.^49^ From the gMSDs, we identify that the *D*_*sp*_ estimated (and consequently *τ*_sat_) at short times show a weak dependency with the number of spike proteins simulated, indicating a diluted condition of spike proteins decorating the envelope (see SI, X.Table IV). Thus, when estimating the lifetime of clusters in Eq. (2), we consider a fixed diffusional time *τ*_sat_ = 90*τ*_sdpd_ regardless of the spike proteins density of the virion. We must highlight, that even though the measured *D*_*sp*_ seems unaffected by the presence of other proteins, the hydrodynamic interactions between spike proteins are effectively influencing the proteins mobility. The plot in FIG. 5.b, shows the probability-density functions (PDFs)— estimated via kernel density estimation (KDE) — of the protein’s squared-displacement (Δ**x**^2^), at three different times (0.1, 10, 90 *τ*_sdpd_). At shorter times, when spike proteins do not interact with others, the distributions remain the same for the different virions. For later times, the number of encounters should be favored for virions with more spike proteins, leading to the small variations observed in the displacements distribution. However, those variations do not affect the *D*_*sp*_ measured (for the number of spike proteins evaluated), as their influence is hindered by the saturation time scale of the gMSD. For virions with a larger number of spike proteins (i.e. Influenza virus^58^ with *N*_*s*_ *>* 400), stronger variations in the displacements distribution would be expected (crowding effects), significantly altering the proteins transport at short time scales, and therefore the effective proteins diffusion *D*_*sp*_.

**FIG. 5.**
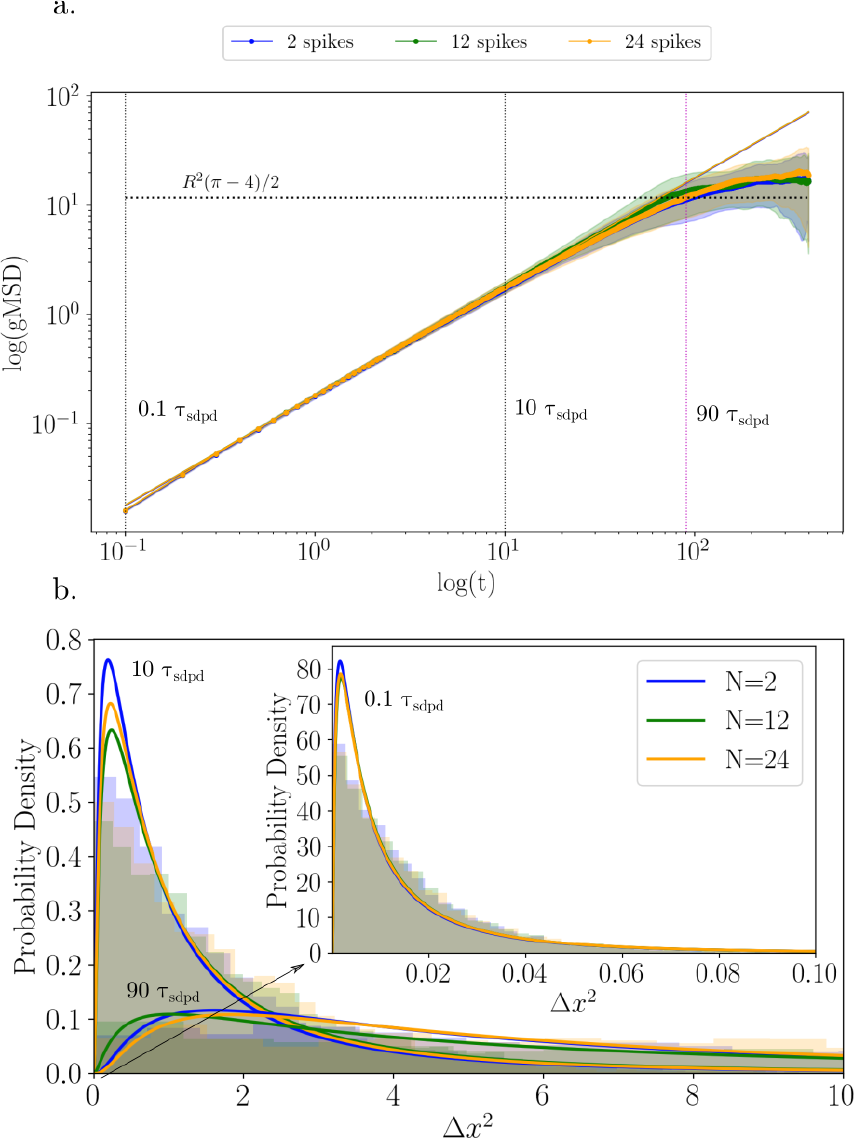
a) gMSD log-log curves for spike protein diffusion on the viral envelope, comparing systems with 2, 12, and 24 spikes proteins. All configurations show similar diffusion dynamics, converging to a plateau due to spatial confinement. The dotted lines indicate threetime points: 0.1*τ*_sdpd_, 10*τ*_sdpd_, and 90*τ*_sdpd_. b) Probability density functions (PDFs) of geodesic distances at Δ*t* = 0.1, 10, and 90*τ*_sdpd_ for systems with 2, 12, and 24 spikes proteins. KDE curves are computed from the average of three replicas (two for the 24-spike case).

In FIG. 6, we now summarize the 3D clustering analysis of spike proteins mediated by hydrodynamic interactions, where we track the number of clusters *N*, cluster size *C*_*s*_, lifetime *l*_*t*_(*C*_*s*_) and lifespan Δ*s*(*C*_*S*_). In FIG. 6.a, we evidence the existence of such clusters, and compare the histogram of *N* for virions with 12 and 24 spike proteins. We observe that virions with 12 spike proteins exhibit *N* values ranging from 5 to 12 clusters, with a mean *N*∼9.5. For 24 spike-proteins case, *N* ranges from 7 to 22, with a mean value *N*∼15. Note that isolated spike proteins are also accounted as clusters of size 1. Thus, in FIG. 6.b, we present the normalized frequencies of the cluster size *C*_*s*_, omitting *C*_*s*_ = 1 to highlight the actual existence of spike proteins aggregates. For both virions, the largest frequencies measured correspond to *C*_*s*_ = 2 and *C*_*s*_ = 3. Even though larger clusters (*C*_*s*_ *>* 3) are also detected, they are significantly less likely for the number of spike-proteins investigated.

**FIG. 6.**
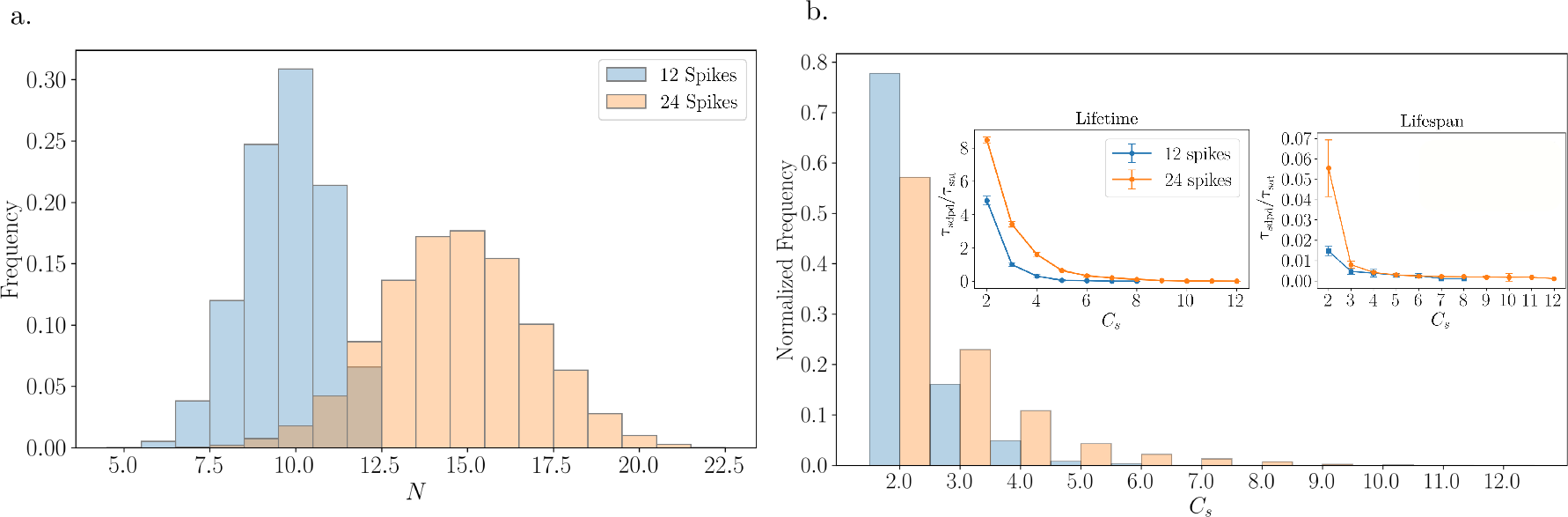
a) Frequency of *N* number of clusters for the 12 and 24 spike proteins simulations. The histogram shows the most common values ranging from 9 to 10 *N* for the 12 spike proteins, while for the 24, the average number is around 15 *N*. b) Normalized frequency for the cluster size *C*_*s*_ for 12 and 24 spike protein simulations. Inside plots are the cluster lifetime [*τ*_sdpd_*/τ*_*sat*_] for different *C*_*s*_ and the lifespan [*τ*_sdpd_*/τ*_*sat*_] per*C*_*s*_ with error bars.

In FIG. 6.b, we further analyze the potential relevance of the formed protein clusters in terms of their *lifetime* Eq.(2) and *lifespan* Eq.(3). Consistent with the cluster size distributions, clusters of size *C*_*s*_ = 2 − 3 amount for largest lifetimes, with few larger clusters (*C*_*s*_ = 4 − 5) exhibiting noticeable lifetimes. However, when analyzing their lifespan,we observe that for cluster size *C*_*s*_ *>* 3 are less stable and to disperse quickly. Interestingly, although clusters of size *C*_*s*_ = 3 exhibit a short measured lifespan, their relatively high frequency suggests that they form frequently despite not persisting over time. Similar clusters containing three proteins have also been reported in the context of HIV,^54,56,57^ raising the question of whether such recurring formations might serve a functional role. However, experimentally determining the lifespan of these clusters remains challenging, making it difficult to assess whether their duration is sufficient to facilitate viral entry. If a longer lifespan is indeed required for functionality, this would suggest that hydrodynamic interactions alone are insufficient to explain their formation. In that case, additional interactions—either between spike proteins themselves or between the proteins and the membrane—may be necessary. Although our model does not explicitly include attractive or repulsive interactions to promote clustering, our results demonstrate that hydrodynamic interactions alone can mediate the formation of spike protein clusters.

Experimentally, the DNA-PAINT imaging technique provides 2D-staggered information about the locations of spike proteins on fixed viral particles, as illustrated in FIG. 7.a for three HIV particle samples (see the full dataset in SI, I.FIG 3). In FIG. 7.a, we present the original DNA-PAINT images (top), the reconstructed protein locations as bright dots (bottom), and the quantified protein clusters. For comparison with simulated virions, we select HIV particles with spike-protein counts in the range 12 ≤ *N*_*sp*_≤ 24. We analyze over 81 viral particles (see SI, I. FIG.4), with envelope and spike-protein radii of *R*∼ 60 nm and *r*_*s*_∼ 6 nm, respectively. Protein clusters are identified using a cutoff radius of 5*r*_*s*_.

**FIG. 7.**
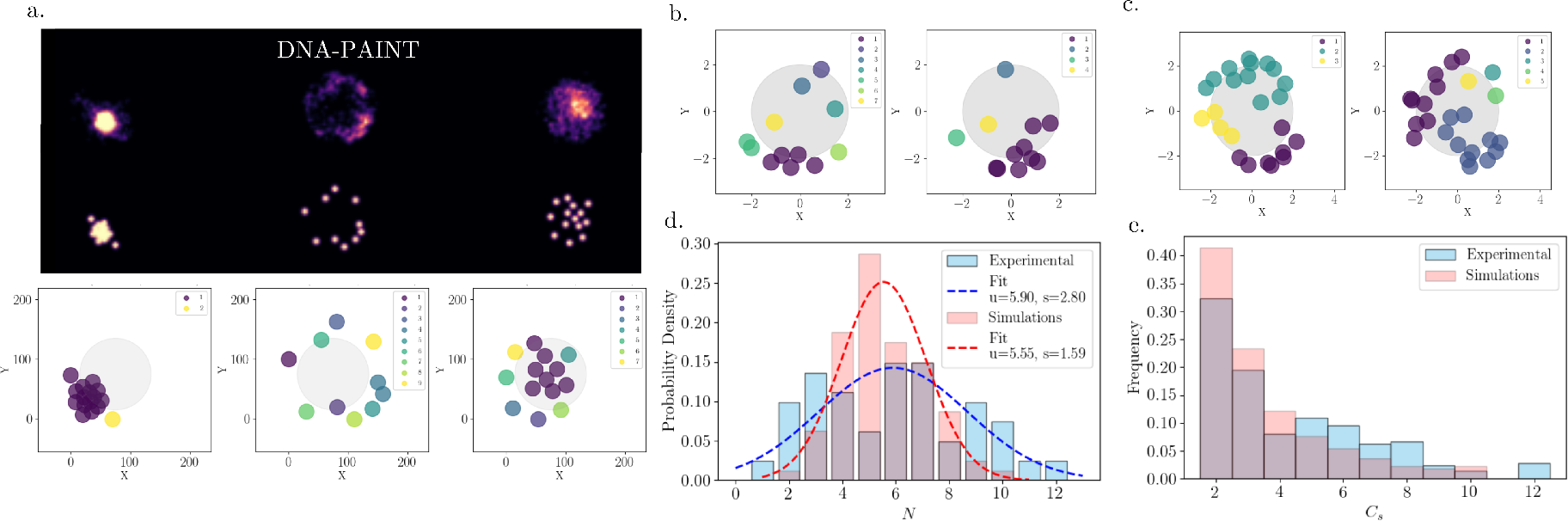
a) Experimental DNA-PAINT images of three viral-particle samples, with their corresponding identified individual proteins location, and clusters. b) and c), the plots show two cases using our simulations for 12 and 24 spike proteins, respectively. The 2D projections illustrate the cluster formation patterns. d) histograms and probability density distributions of *N*, for HIV-1 viral particles with 12 ≤ *N*_*sp*_ ≤ 24 and the combined simulations of virion models with 12 and 24 spike proteins. e) Clusters size *C*_*s*_ distributions in both experiments and simulations, indicating a predominant formation of doublets and triplets.

To facilitate comparison with simulations, we perform 2D clustering analysis on simulated virions, combining data sets of systems with 12 and 24 spike proteins. FIG. 7.b and FIG. 7.c illustrate the clustering results for simulated virions with 12 and 24 spike proteins, respectively. We select 80 frames from each simulation set (see SI, I. FIG 5), spaced over time intervals *t >* 10*τ*_*sat*_, ensuring that the frames are uncorrelated in time to match the diversity of the experimental measurements.

Overall, clustering patterns are qualitatively similar (see SI, I. FIG 4, 5) in both experimental and simulated data. Quantitatively, both exhibit similar distributions in number and size of the clusters. In FIG. 7.d, we compare the distribution of the number of clusters *N* between experiments and simulations. The fitted Gaussian probability distribution functions show similar peaks at *N* = 6 for both cases. However, the standard deviation is larger for the experimental data (*s* = 2.8) than for the simulations (*s* = 1.59), indicating a broader distribution in the number of clusters observed experimentally, likely associated to the presence of clusters of larger size over the different samples.

In particular, compact and large clusters are more prevalent in the experimental data, while in simulations scattered small protein clusters are dominant. In FIG. 7.e, we compare the distribution of clusters size (omitting *C*_*s*_ = 1), showing a predominant population of doublets *C*_*s*_ = 2 and triplets *C*_*s*_ = 3, in both experiments and simulations. However, the formation of clusters *C*_*s*_ *>* 5 are favored in experimental observations. This suggests that whereas hydrodynamic interactions are sufficient to stabilize small clusters, additional interactions, such as those mediated by the viral envelope or enhanced protein-protein interactions (i.e. HIV-1 proteins-charge distribution), may play a crucial role in the formation of larger clusters. Further improvements in the simulation model, incorporating additional spike-protein interactions and explicit drag forces with the viral envelope, will be addressed in future work to reproduce experimental observations in HIV-1 virions with different degrees of maturation.

## IV. CONCLUSIONS

In this work, we investigated how spike flexibility influences the passive transport of viruses. Adopting mesoscalerigid-body descriptions, that incorporate spike flexibility features, together with a fluctuating hydrodynamics via the SDPD method, allows us to model more realistic virion dynamics on temporal and spatial scale of the whole virion. Additionally it offers an improved representation to investigate and understand viral transport and entry.

Our results show that incorporating structural flexibility into virus models leads to slight changes in rotational mobility compared to recently proposed rigid models. Interestingly, while flexibility does affect rotational motion, it does not significantly alter translational diffusion across the models we studied. Overall, we find that the viral envelope, due to its larger size, primarily determines the characteristic timescales of passive virion transport. Nonetheless, the dynamic behavior of spike proteins continues to impact viral mobility, despite their smaller size and faster timescales. This underscores the interplay between the motion of the envelope and the spike proteins in shaping the overall transport properties of the viruses.

The experimental results using DNA-PAINT super-resolution imaging of HIV-1 particles provides a valuable experimental reference to qualitatively assess the clustering behavior observed in our simulations. Remarkably, the clustering behavior of spike proteins (particularly the frequent formation of transient triplets) mediated by hydrodynamic interactions poses the question about the potential functional role in viral infectivity. These simulation-based observations, which qualitatively align with reports from HIV studies,^10,54,56,57^ point to a possible correlation between clustering properties, such as lifespan and aggregation probability, and the virus’s ability to engage host cells. Although direct time-resolved experimental validation of real virions is currently limited, our results underscore the power of computational modeling to probe complex, time-dependent interactions that are difficult to capture in vitro. More broadly, the SDPD methodology introduced here provides a flexible framework for investigating nanoscale biological systems with similar structural and dynamical characteristics, extending beyond the specific case of virion transport.

## SUPPLEMENTARY MATERIAL

The supporting material includes additional information relevant to the simulations presented in this study. It contains detailed simulation parameters, time scales, resolution tests, angle parameters, correction factors for diffusion coefficients, and squared metrics such as MSAD, MSD, and gMSD.

### ACKNOWLEDGMENTS

D.M.C, F.B.U, M.E and N.M acknowledge funding by the Basque Government through the BERC 2022-2025 program and by the Ministry of Science, Innovation and Universities: BCAM Severo Ochoa accreditation CEX2021-001142S/MICIN/AEI/10.13039/501100011033. The Spanish State Research Agency through the project PID2020-117080RBC55 funded by (AEI/FEDER, UE) with acronym COMPUNANO-HYDRO. The computer resources at MareNostrum and the technical support provided by BSC (RES-FI2024-3-0014 Mesoscale Modelling of Passive Transport of Viruses and Viral Spikes). C.Z. acknowledges the Human Frontier Science Program Organization (HFSP) through a cross-disciplinary post-doctoral fellowship (LT0025/2023-C),H.S.H. the Biotechnology and Biological Sciences Research Council (BBSCR) through the London Interdisciplinary Doctoral Programme (BB/XXXXXX), and S.S the Royal Society through a Dorothy Hodgkin fellowship (DHF R1 191019). This work has also been supported by The Chan Zuckerberg Initiative “Multi-color single molecule tracking with lifetime imaging” (2023-321188). S.P.P acknowledges funding from the European Research Council [ERC-2019-CoG863869 FUSION]

## V. AUTHOR CONTRIBUTIONS

D.M and N.M conceived the numerical and modeling study. The first draft of the manuscript was written by N.M and D.MC. D.MC conducted the simulations and data processing. D.M, F.B.U, M.E and N.M discussed and analyzed the numerical results. C.Z, H.S.H, D.W, C.I, S.S and S.P.P contribute with the experimental results on DNA-PAINT superresolution imaging of HIV-1 particles, production and functional validation of labeled HIV-1 viruses. All authors discussed the results and approved the final version of the manuscript.

## DATA AVAILABILITY STATEMENT

The data that support the findings of this study are available from the corresponding author upon reasonable request.

## References

1 A. Valero-Rello, C. Baeza-Delgado, I. Andreu-Moreno, and R. Sanjuán, “Cellular receptors for mammalian viruses,” PLoS Pathogens 20, e1012021 (2024).

2 B. Turoňová, M. Sikora, C. Schürmann, W. J. Hagen, S. Welsch, F. E. Blanc, S. von Bülow, M. Gecht, K. Bagola, and C. Hörner, “In situ structural analysis of sars-cov-2 spike reveals flexibility mediated by three hinges,” Science 370, 203–208 (2020).

3 L. Casalino, C. Seitz, J. Lederhofer, Y. Tsybovsky, I. A. Wilson, M. Kanekiyo, and R. E. Amaro, “Breathing and tilting: mesoscale simulations illuminate influenza glycoprotein vulnerabilities,” ACS central science 8, 1646–1663 (2022).

4 D. Chmielewski, E. A. Wilson, G. Pintilie, P. Zhao, M. Chen, M. F. Schmid, G. Simmons, L. Wells, J. Jin, A. Singharoy, et al., “Structural insights into the modulation of coronavirus spike tilting and infectivity by hinge glycans,” Nature Communications 14, 7175 (2023).

5 T. Sakai, H. Takagi, Y. Muraki, and M. Saito, “Unique directional motility of influenza c virus controlled by its filamentous morphology and shortrange motions,” Journal of virology 92, 10–1128 (2018).

6 M. D. Vahey and D. A. Fletcher, “Influenza a virus surface proteins are organized to help penetrate host mucus,” Elife 8, e43764 (2019).

7 L. Stevens, S. de Buyl, and B. M. Mognetti, “The sliding motility of the bacilliform virions of influenza a viruses,” Soft matter 19, 4491–4501 (2023).

8 S. Yang, G. Hiotis, Y. Wang, J. Chen, J.-h. Wang, M. Kim, E. L. Reinherz, and T. Walz, “Dynamic hiv-1 spike motion creates vulnerability for its membrane-bound tripod to antibody attack,” Nature Communications 13, 6393 (2022).

9 L. Andronov, M. Han, Y. Zhu, A. Balaji, A. R. Roy, A. E. Barentine, P. Patel, J. Garhyan, L. S. Qi, and W. E. Moerner, “Nanoscale cellular organization of viral rna and proteins in sars-cov-2 replication organelles,” Nature Communications 15 (2024).

10 I. C. Andres, T. Malinauskas, and S. Padilla-Parra, “Structure and dynamics of hiv-1 env trimers on native virions engaged in living t cells,” Biophysical Journal 120, 130a (2021).

11 H. Gelderblom, M. Özel, and G. Pauli, “Morphogenesis and morphology of hiv structure-function relations,” Archives of virology 106, 1–13 (1989).

12 J. Chojnacki, D. Waithe, P. Carravilla, N. Huarte, S. Galiani, J. Enderlein, and C. Eggeling, “Envelope glycoprotein mobility on HIV-1 particles depends on the virus maturation state,” Nature Communications 8 (2017).

13 Y. G. Kuznetsov and A. McPherson, “Atomic force microscopy in imaging of viruses and virus-infected cells,” Microbiology and Molecular Biology Reviews 75, 268–285 (2011).

14 A. Toshio, U. Takayuki, and F. Takeshi, “High-speed atomic force microscopy for nano-visualization of dynamic biomolecular processes,” Progress in Surface Science 83, 337–437 (2008).

15 W. Godinez, L.M. S. Wörz, B. Müller, R. Eils, and K. Rohr, “Deterministic and probabilistic approaches for tracking virus particles in timelapse fluorescence microscopy image sequences,” Medical Image Analysis 13, 325–342 (2009), includes Special Section on Functional Imaging and Modelling of the Heart.

16 R. Gordon, “Biosensing with nanoaperture optical tweezers,” Optics and Laser Technology 109, 328–335 (2019).

17 C. J. Bustamante, Y. R. Chemla, S. Liu, and M. D. Wang, “Optical tweezers in single-molecule biophysics,” Nature Reviews Methods Primers 1, 25 (2021).

18 G. Katz, Y. Benkarroum, H. Wei, W. J. Rice, D. Bucher, A. Alimova,A. Katz, J. Klukowska, G. T. Herman, and P. Gottlieb, “Morphology of influenza b/lee/40 determined by cryo-electron microscopy,” PLOS ONE 9, 1–8 (2014).

19 Z. Ke, J. Oton, K. Qu, M. Cortese, V. Zila, L. McKeane, T. Nakane, J. Zivanov, C. J. Neufeldt, B. Cerikan, J. M. Lu, J. Peukes, X. Xiong, H. G. Kräusslich, S. H. Scheres, R. Bartenschlager, and J. A. Briggs, “Structures and distributions of SARS-CoV-2 spike proteins on intact virions,” Nature 588, 498–502 (2020).

20 A. Slater, N. Nair, R. Suétt, R. Mac Donnchadha, C. Bamford, S. Jasim, D. Livingstone, and E. Hutchinson, “Visualising viruses,” Journal of General Virology 103, 001730 (2022).

21 E. E. Jefferys and M. S. P. Sansom, “Chapter 10: Computational virology: Molecular simulations of virus dynamics and interactions,” Physical Virology Advances in Experimental Medicine and Biology, 1215 (2019).

22 B. Subhomoi, D. Debajit, H. Zaved, J. Amit, and T. Keshawanand, “Unravelling viral dynamics through molecular dynamics simulations - a brief overview,” Biophysical Chemistry 291, 106908 (2022).

23 H. Ode, M. Nakashima, S. Kitamura, W. Sugiura, and H. Sato, “Molecular dynamics simulation in virus research,” Frontiers in microbiology 3, 258 (2012).

24 M. R. Machado and S. Pantano, “Fighting viruses with computers, right now,” Current Opinion in Virology 48, 91–99 (2021).

25 P. E. Jones, C. Pérez-Segura, A. J. Bryer, J. R. Perilla, and J. A. Hadden-Perilla, “Molecular dynamics of the viral life cycle: progress and prospects,” Current Opinion in Virology 50, 128–138 (2021).

26 D. L. Lynch, A. Pavlova, Z. Fan, and J. C. Gumbart, “Understanding virus structure and dynamics through molecular simulations,” Journal of chemical theory and computation 19, 3025–3036 (2023).

27 Y. Li, Y.-L. Zhu, Y.-C. Li, H.-J. Qian, and C.-C. Sun, “Self-assembly of two-patch particles in solution: A brownian dynamics simulation study,” Molecular Simulation 40, 449–457 (2014).

28 M. A. Islam, S. Barua, and D. Barua, “A multiscale modeling study of particle size effects on the tissue penetration efficacy of drug-delivery nanoparticles,” BMC Systems Biology 11 (2017).

29 J. Ilnytskyi, “Self-Assembly of Nanoparticles Decorated by Liquid Crystalline Groups: Computer Simulations,” Self-Assembly of Nanostructures and Patchy Nanoparticles (2020).

30 V. E. Debets, L. M. C. Janssen, and A. S. Saric, “Characterising the diffusion of biological nanoparticles on fluid and cross-linked membranes,” Soft Matter 16, 10628 (2020).

31 J. Liu, R. Tourdot, V. Ramanan, N. J. Agrawal, and R. Radhakrishanan, “Mesoscale simulations of curvature-inducing protein partitioning on lipid bilayer membranes in the presence of mean curvature fields,” Molecular Physics 110, 1127–1137 (2012).

32 N. Moreno, B. Sutisna, and E. Fried, “Entropic factors and structural motifs of triblock-terpolymer-based patchy nanoparticles,” Royal Society of Chemistry (2020).

33 S. B. Chen, “Dissipative Particle Dynamics Simulation of Nanoparticle Diffusion in a Crosslinked Polymer Network,” The Journal of Physical Chemistry B (2022).

34 L. Li, X. Li, Z. Duan, R. J. Meyer, R. Carr, S. Raman, L. Koziol, and G. Henkelman, “Adaptive kinetic Monte Carlo simulations of surface segregation in PdAu nanoparticles,” Nanoscale 11, 10524–10535 (2019).

35 M. A. Kanso, J. H. Piette, J. A. Hanna, and A. J. Giacomin, “Coronavirus rotational diffusivity,” Physics of Fluids 32 (2020).

36 M. A. Kanso, V. Chaurasia, E. Fried, and A. J. Giacomin, “Peplomer bulb shape and coronavirus rotational diffusivity,” Physics of Fluids 33 (2021).

37 F. Balboa Usabiaga and B. Delmotte, “A numerical method for suspensions of articulated bodies in viscous flows,” Journal of Computational Physics 464, 111365 (2022).

38 N. Moreno, D. M. Chaparro, and F. B. Usabiaga, “Hydrodynamics of spike proteins dictate a transport - affinity competition for SARS - CoV - 2 and other enveloped viruses,” Scientific Reports, 1–13 (2022).

39 D. Moreno-Chaparro, N. Moreno, F. B. Usabiaga, and M. Ellero, “Computational modeling of passive transport of functionalized nanoparticles,” The Journal of Chemical Physics 158, 104108 (2023).

40 P. Espanol and M. Revenga, “Smoothed dissipative particle dynamics,” Phys. Rev. E 67, 026705 (2003).

41 M. Ellero and P. Espanol, “Everything you always wanted to know about SDPD (but were afraid to ask),” Appl. Math. Mech.-Engl. Ed 39, 103–124 (2018).

42 J. J. Monaghan, “Smoothed particle hydrodynamics,” In: Annual review of astronomy and astrophysics 30, 543–574 (1992).

43 N. Moreno and M. Ellero, “Arbitrary flow boundary conditions in smoothed dissipative particle dynamics: A generalized virtual rheometer,” Physics of Fluids 33, 012006 (2021).

44 E. Zohravi, N. Moreno, and M. Ellero, “Computational mesoscale framework for biological clustering and fractal aggregation,” Soft Matter 19, 7399–7411 (2023).

45 E. Zohravi, N. Moreno, K. Hawkins, D. Curtis, and M. Ellero, “Mesoscale modelling of fibrin clots: the interplay between rheology and microstructure at the gel point,” Soft Matter 21, 1141–1151 (2025).

46 S. Espinosa-Moreno, N. Moreno, and M. Ellero, “Computational modelling of thixotropic multiphase fluids using smoothed dissipative particle dynamics,” 10.48550/arXiv.2507.02766 (2025).

47 A. Vázquez-Quesada and M. Ellero, “GENERIC-compliant simulations of Brownian multi-particle systems: modeling stochastic lubrication,” SeMA Journal 79, 165–185 (2022).

48 A. Ledesma-Durán, J. Munguía-Valadez, J. A. Moreno-Razo, S.I. Hernández, and I. Santamaría-Holek, “Entropic Effects of Interacting Particles Diffusing on Spherical Surfaces,” Frontiers in Physics 9 (2021).

49 A. Ledesma-Durán and L.H. Juárez-Valencia, “Diffusion coefficients and MSD measurements on curved membranes and porous media,” European Physical Journal E 46 (2023).

50 J. Padding and A. Louis, “Hydrodynamic interactions and brownian forces in colloidal suspensions: Coarse-graining over time and length scales,” Physical Review E-Statistical, Nonlinear, and Soft Matter Physics 74, 031402 (2006).

51 X. Bian, S. Litvinov, R. Qian, M. Ellero, and N. Adams, “Multiscale modeling of particle in suspension with smoothed dissipative particle dynamics,” Physics of Fluids 24 (2012).

52 A. Einstein, “On the movement of small particles suspended in stationary liquids required by the molecular,” Ann. d. Phys (1905).

53 J. Schnitzbauer, M. T. Strauss, T. Schlichthaerle, F. Schueder, and R. Jungmann, “Super-resolution microscopy with dna-paint,” Nature protocols 12, 1198–1228 (2017).

54 D. J. Williamson, C. Zaza, I. Carlon-Andres, T. Starling, A. Gentili, J. W. Thrush, A. Le Bas, R. T. Ravi, S. Neil, R. J. Owens, M. Dumoux, S. Simoncelli, and S. Padilla-Parra, “Single-molecule localisation microscopy approaches reveal envelope glycoprotein clusters in single-enveloped viruses: a potential functional role?” Biochemical Society Transactions, BST20240769 (2025).

55 M. D. Joseph, E. Tomas Bort, R. P. Grose, P. J. McCormick, and S. Simoncelli, “Quantitative super-resolution imaging for the analysis of gpcr oligomerization,” Biomolecules 11, 1503 (2021).

56 J. Liu, A. Bartesaghi, M. J. Borgnia, G. Sapiro, and S. Subramaniam, “Molecular architecture of native HIV-1 gp120 trimers,” Nature 455, 109– 113 (2008).

57 B. K. Ganser-Pornillos, M. Yeager, and O. Pornillos, “Assembly and architecture of HIV,” Advances in Experimental Medicine and Biology 726, 441–465 (2012).

58 A. Harris, G. Cardone, D. C. Winkler, J. B. Heymann, M. Brecher, J. M. White, and A. C. Steven, “Influenza virus pleiomorphy characterized by cryoelectron tomography,” Proceedings of the National Academy of Sciences of the United States of America 103, 19123–19127 (2006).

