## Supplementary Information for "Mesoscale Transport of Enveloped Viruses"

The supporting material includes additional information relevant to the simulations presented in this study. It contains detailed simulation parameters, time scales, resolution tests, angle parameters, correction factors for diffusion coefficients, and metrics such as MSAD, MSD, and gMSD.

### I. SIMULATIONS PARAMETERS AND RELEVANT TIME SCALES

To characterize the spike proteins and the dynamics of the whole virion, we can identify different characteristic time scales, such as the sonic time, kinematic time, and Brownian time. The sonic time is given by $t_{cs} = R_s/c_s$, where $R_s$ is the spike or envelope radius and $c_s$ is the speed of sound. The kinematic time is $\tau_\nu = R_s^2/\nu$, where $\nu = \eta/\rho$ is the kinematic viscosity, and the Brownian time is $\tau_D = R_s^2/D$, with $D$ the diffusion coefficient.

Taking into account that these time scales must satisfy the inequality $t_{cs} < \tau_\nu < \tau_D$ [1], we set the thermal energy of the system as $k_B T = 1[\varepsilon_{\text{sdpd}}]$, the speed of sound $c_s = 40[l_{\text{sdpd}}/\tau_{\text{sdpd}}]$ and fluid viscosity $\eta = 5[m_{\text{sdpd}} l_{\text{sdpd}}/\tau_{\text{sdpd}}]$. The radius of each SDPD particle is $a = 0.1 l_{\text{sdpd}}$ the particle mass is $m = 0.008 m_{\text{sdpd}}$ and mass density $\rho_a = 1.9[m_{\text{sdpd}}/l_{\text{sdpd}}^2]$, yielding a thermal velocity $v_{th} = \sqrt{k_B T/m} = 11.2[l_{\text{sdpd}}/\tau_{\text{sdpd}}]$.

Similarly, for the envelope constructed with 3993 SDPD particles, the mass is $M = 3993m[m_{\text{sdpd}}]$, the mass density is $\rho_0 = M/V_e = 0.95[m_{\text{sdpd}}/l_{\text{sdpd}}^3]$, where $V_e = 4/3\pi R^3[l_{\text{sdpd}}^3]$ is the envelope volume. Finally, the thermal velocity of a envelope is $V_{th} = \sqrt{k_B T/M} = 0.177[l_{\text{sdpd}}/\tau_{\text{sdpd}}]$ Using these parameters, we compute the relevant time scales for the spike proteins with size $r_s = 0.24 l_{\text{sdpd}}$, we have $t_{cs} = 0.006 < \tau_\nu = 0.011 < \tau_D = 1.302$, whereas for the envelope with $R = 2l_{\text{sdpd}}$, we have $t_{cs} = 0.05 < \tau_\nu = 0.80 < \tau_D = 753.9$. In Table I, we give a breakdown of the key geometrical, physical, and simulation parameters used.

Additionally, we include other relevant dimensionless numbers that characterize the fluid regime in our simulations, such as Schmidt number $\text{Sc} = \nu/D$, Peclet number $\text{Pe} = V_{th} R_s/D$, and the Reynolds number $\text{Re} = \rho_0 V_{th}(2R_s)/\eta$ computed using the thermal velocity. To ensure numerical stability and lower density fluctuation, we use a time step size $dt = 0.001[\tau_{\text{sdpd}}]$, taking into account the criteria $dt < min(0.25h/c_s, 0.125h^2/\nu)$. [2]

TABLE I. List of key parameters and dimensionless numbers. The length units are $l_{\text{sdpd}}$, time units are $\tau_{\text{sdpd}}$, and mass units are $m_{\text{sdpd}}$.

| **Parameter** | **Value** | **Units** |
|---|---|---|
| **Geometrical** | | |
| Interparticle distance ($dx$) | 0.2 | $l_{\text{sdpd}}$ |
| SDPD particle radius ($a$) | 0.1 | $l_{\text{sdpd}}$ |
| Particle density ($d$) | $1/dx^3$ | $l_{\text{sdpd}}^{-3}$ |
| Radius of envelope ($R/dx$) | 10 | - |
| Radius of spike ($r_s/dx$) | 1 | - |
| Interaction radius ($h/dx$) | 4 | - |
| **Physical Properties** | | |
| Density ($\rho_0$) | 0.95 | $m_{\text{sdpd}}/l_{\text{sdpd}}^3$ |
| Mass particle ($m$) | 0.008 | $m_{\text{sdpd}}$ |
| Mass envelope ($M$) | 31.9 | $m_{\text{sdpd}}$ |
| Thermal energy ($K_bT$) | 1 | $\varepsilon_{\text{sdpd}}$ |
| Viscosity ($\eta$) | 5 | $m_{\text{sdpd}}/(l_{\text{sdpd}}\tau_{\text{sdpd}})$ |
| Speed of sound ($c$) | 40 | $l_{\text{sdpd}}/\tau_{\text{sdpd}}$ |
| Thermal speed particle ($v_{th}$) | 11.2 | $l_{\text{sdpd}}/\tau_{\text{sdpd}}$ |
| Thermal speed envelope ($V_{th}$) | 0.17 | $l_{\text{sdpd}}/\tau_{\text{sdpd}}$ |
| **Dimensionless Numbers** | | |
| Reynolds number (Re) | 0.13 | |
| Schmidt number (Sc) | 988.7 | |
| Peclet number (Pe) | 133.4 | |
| **Time Parameters** | **Envelope** | **Spike** |
| Time step ($dt$) | $1e^{-3}$ | $1e^{-4}$ |
| Diffusive time ($\tau_D$) | 753.9 | 1.302 |
| Kinematic time ($\tau_v$) | 0.8 | 0.011 |
| Sonic time ($\tau_{cs}$) | 0.05 | 0.006 |
| Ratio $\tau_D/\tau_v$ | 942.5 | 113.1 |

### II. RESOLUTION STUDY

To ensure the accuracy of our simulations, we conduct a resolution study to determine the minimal spatial discretization required for reliable results. To determine the minimal resolution, we first estimate the spatial discretization that provides an adequate estimation of the rotational mobility ($M_0 = 1/(8\pi\eta R^3)$) of an spherical object in athermal conditions $k_B T = 0$. We focus on the rotational mobility as it is more sensitive to spatial discretization than translational mobility, due to its dependence on $R^3$. We simulate spherical envelopes for $R/dx$ values in the range $\{5, 10, 15\}$, with its center of mass fixed, and apply an external torque $\mathscr{T}$ along the $z$-axis, inducing a steady angular velocity $\omega_z$ in the $xy$-plane. To ensure consistency, the Reynolds number is kept as $\mathrm{Re} \ll 1$ in all simulations. The numerical mobility is calculated as $M = \omega_z/\mathscr{T}$, and the ratio $M/M_0$ is used to identify the resolution that balances computational efficiency and accuracy. We identify that a ratio of $R/dx = 10$ provides a sufficient approximation of the rotational mobility, yielding a mobility ratio $M/M_0 \approx 1.05$.

After identifying this optimal resolution, we compute the rotational diffusion coefficient $D_r^e$ and the translational diffusion coefficient $D_t^e$ of an spherical envelope freely moving in a Newtonian fluid, using their mean square displacement (MSD) and mean square angular displacement (MSAD). We perform three independent realizations to ensure statistical robustness. The MSAD curve exhibits an early linear region before reaching a plateau, which is consistent with theoretical predictions, where this saturation occurs near $(\pi^2 - 4)/2$ [3]. FIG. 1 presents the MSD and MSAD curves, averaged over the three realizations. We highlight the fitting intervals used for diffusion estimation, from 10 to $100\tau_\nu$ for rotational and translational diffusion. After applying the correction in the measured diffusion due to periodic boundary conditions as described in SI.V. , we estimate the relative errors of the diffusion coefficients with respect to the theoretical Stokes-Einstein relation, on the order of 6% and 24%, for $D_t$ and $D_r$, respectively. Larger errors in rotational diffusion are associated with its $R^3$ dependence, which amplifies the impact of finite-size effects.

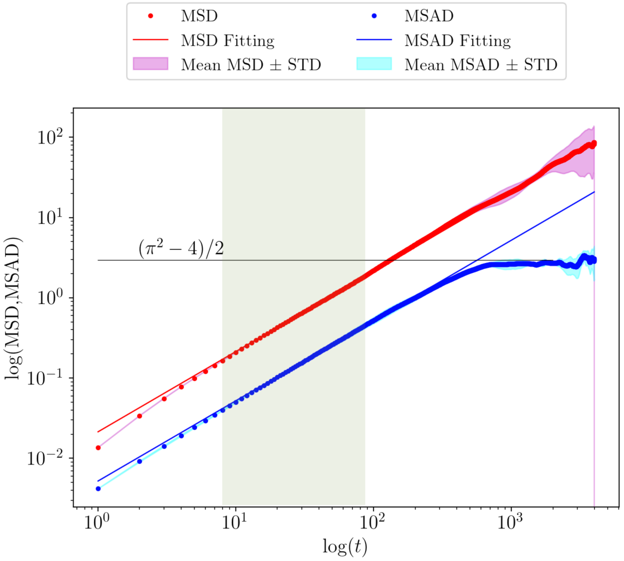


FIG. 1. Mean square displacement MSD and mean square angular displacement MSAD log-log curves for a spherical envelope under three different random fluid conditions. The shaded green region (10 to 100$\tau_v$) marks the time interval used to estimate the rotational diffusion coefficient $D_r^e$ and the translational diffusion coefficient $D_t^e$. The MSAD curve saturates near the theoretical limit $(\pi^2 - 4)/2$

### III. THEORETICAL TILT ANGLE VARIATION

TABLE II. Theoretical mean tilt angles $\beta$ for different angular stiffness values $K_\alpha$.

| $K_\alpha$ | **Mean angle $\beta$ (°)** | $K_\alpha$ | **Mean angle $\beta$ (°)** |
|---|---|---|---|
| 0 | 90.00 | 50 | 8.07 |
| 1 | 49.29 | 100 | 5.72 |
| 5 | 24.80 | 500 | 2.56 |
| 10 | 17.82 | 1000 | 1.81 |

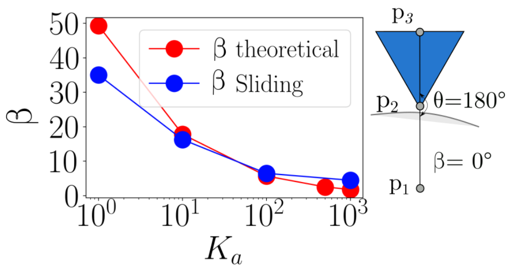


FIG. 2. Tilting angle $\beta$ as a function of angular stiffness $K_\alpha$ (1 to 1000) for both theoretical predictions and simulation results on curved (envelope) surfaces. $\beta$ is defined between three points: envelope center of mass ($p_1$), spike base ($p_2$), and spike tip ($p_3$). Higher $K_\alpha$ values restrict spike motion, reducing $\beta$.

### IV. DIFFUSION COEFFICIENTS VARIATIONS WITH $K_\alpha$

TABLE III. Normalized diffusion coefficients ($\bar{D}_t$ and $\bar{D}_r$) with standard deviations for varying $K_\alpha$ values.

| $K_\alpha$ | $\bar{D}_t \pm \sigma$ | $\bar{D}_r \pm \sigma$ |
|---|---|---|
| 10 | $0.972 \pm 0.002$ | $0.947 \pm 0.031$ |
| 100 | $0.944 \pm 0.059$ | $0.924 \pm 0.055$ |
| 1000 | $1.018 \pm 0.031$ | $0.905 \pm 0.045$ |

### V. CORRECTION FACTOR FOR DIFFUSION COEFFICIENTS

To accurately compare simulated diffusion coefficients with theoretical predictions, correction factors are applied to account for finite-size effects in the simulation box.

#### A. Volume Fraction Calculation

The particle volume fraction $\phi$ is computed using the particle radius $r$ and the cubic box length $L$:

$$\phi = \frac{4\pi R^3}{3L^3}$$

For this analysis, the parameters are:

$$R = 2, \quad L = 15$$

#### B. Translational Diffusion Correction Factor

The correction factor $\gamma$ [1, 4] for the translational diffusion coefficient $D_t$ is calculated using an empirical formula:

$$a = 1.7601\phi^{1/3}, \quad b = 1.5593\phi^2,$$

$$c = 3.9799\phi^{8/3}, \quad d = 3.0734\phi^{10/3}$$

$$\gamma = 1 - a + \phi - b + c - d$$

The corrected translational diffusion coefficient is given by:

$$D_t = D_t^o \gamma = \frac{\gamma}{6\pi\eta R}$$

where $D_t^o$ is the classical Stokes-Einstein coefficient, $\eta = 5$ is the fluid viscosity, and $R$ is the hydrodynamic (envelope) radius.

#### C. Rotational Diffusion Correction Factor

A correction factor $C$ is applied to the rotational diffusion coefficient [5]. The corrected rotational diffusion is given by:

$$C = \frac{1}{6\eta L^3}$$

$$D_r = D_r^o - C = \left(\frac{1}{8\pi\eta R^3}\right) - C$$

where $D_r^o$ is the classical Stokes-Einstein expression for rotational diffusion.

In this case, the correction factor $C$ is less significant than for the translational case.

#### D. Comparison Between Simulated and Corrected Diffusion

The simulated diffusion coefficients $D_t^{\text{simul}}$ and $D_r^{\text{simul}}$ are normalized by their respective corrected values:

$$\frac{D_t^{\text{simul}}}{D_t}, \quad \frac{D_r^{\text{simul}}}{D_r}$$

These ratios provide insight into how closely the simulation results match theoretical expectations, accounting for both system size and viscosity.

### VI. MEAN SQUARED ANGULAR DISPLACEMENT (MSAD)

The Mean Squared Angular Displacement (MSAD) quantifies the rotational diffusion of particles using spherical coordinates derived from 3D Cartesian data.

To compute angular displacement, Cartesian coordinates $(x, y, z)$ are first converted to spherical coordinates $(r, \theta, \phi)$ using the following formulas:

$$r = \sqrt{x^2 + y^2 + z^2}$$

$$\theta = \cos^{-1}\left(\frac{z}{r}\right) \quad \text{if } r \neq 0$$

$$\phi = \tan^{-1}(y, x)$$

If $r = 0$, both $\theta$ and $\phi$ are set to 0.

The MSAD is computed for each lag time $\Delta t$ as follows:

$$\Delta\alpha^2 = \left[\cos^{-1}\left(\sin(\phi_1)\sin(\phi_2) + \cos(\phi_1)\cos(\phi_2)\cos(\theta_1 - \theta_2)\right)\right]^2$$

.

Where $\phi_1 = \phi(t + \Delta t)$, $\phi_2 = \phi(t)$, $\phi_1 = \theta_1(t + \Delta t)$ and $\theta_2 = \phi(t)$

$$\text{MSAD}(\Delta t) = \frac{1}{N - \Delta t} \sum_{t=1}^{N-\Delta t} \Delta\alpha^2(\Delta t)$$

**Diffusion coefficient estimation:** From MSAD curve, the rotational diffusion coefficient $D_r$ is computed using the following equation:

$$\text{MSAD}(\Delta t) = 4D_r t \Rightarrow \quad D_r = \frac{\text{MSAD}(t)}{4t}$$

### VII. MEAN SQUARED DISPLACEMENT (MSD)

The Mean Squared Displacement (MSD) is a widely used metric for characterizing translational diffusion from particle trajectory data in 3D space.

For each lag time $\Delta t$, the displacement is computed in each spatial dimension:

$$\Delta x_i(\Delta t) = x(t_i + \Delta t) - x(t_i)$$

$$\Delta y_i(\Delta t) = y(t_i + \Delta t) - y(t_i)$$

$$\Delta z_i(\Delta t) = z(t_i + \Delta t) - z(t_i)$$

The total squared displacement is:

$$\Delta r_i^2(\Delta t) = (\Delta x_i)^2 + (\Delta y_i)^2 + (\Delta z_i)^2$$

The MSD is obtained by averaging over all valid starting points:

$$\mathrm{MSD}(\Delta t) = \frac{1}{N - \Delta t} \sum_{i=1}^{N-\Delta t} \Delta r_i^2(\Delta t)$$

MSD can be decomposed into its directional components:

$$\mathrm{MSD}_x(\Delta t) = \frac{1}{N - \Delta t} \sum_{i=1}^{N-\Delta t} (\Delta x_i)^2$$

$$\mathrm{MSD}_y(\Delta t) = \frac{1}{N - \Delta t} \sum_{i=1}^{N-\Delta t} (\Delta y_i)^2$$

$$\mathrm{MSD}_z(\Delta t) = \frac{1}{N - \Delta t} \sum_{i=1}^{N-\Delta t} (\Delta z_i)^2$$

**Diffusion Coefficient Estimation:** From the MSD curve, the translational diffusion coefficient $D_t$ is extracted using theoretical formulations:

$$\mathrm{MSD}(t) = 6D_t t \quad \Rightarrow \quad D_t = \frac{\mathrm{MSD}(t)}{6t}$$

### VIII. GEODESIC MEAN SQUARED DISPLACEMENT (GMSD)

The geodesic MSD (gMSD) [3, 6] accounts for angular distances on a spherical surface and is useful for confined diffusion on curved surfaces.

The gMSD is computed using the spherical law of cosines multiplied by the envelope radius $R$:

$$\mathrm{gMSD}(\Delta t) = \frac{1}{N - \Delta t} \sum_{i=1}^{N-\Delta t} \Delta d^2(\Delta t)$$

$$\Delta d^2 = \left[R \cos^{-1}\left(\sin(\phi_1)\sin(\phi_2) + \cos(\phi_1)\cos(\phi_2)\cos(\theta_1 - \theta_2)\right)\right]^2$$

However, we convert them to Cartesian coordinates on the unit sphere using unit vectors to compute the angular displacement,

$$\vec{u}(\theta,\phi) = \begin{bmatrix} x \\ y \\ z \end{bmatrix} = \begin{bmatrix} \sin(\theta)\cos(\phi) \\ \sin(\theta)\sin(\phi) \\ \cos(\theta) \end{bmatrix}.$$

These vectors describe the direction of each point.

We compute the angular displacement by computing angle $\alpha_i$ between $\vec{u}_i$ and $\vec{u}_{i+\Delta t}$ that corresponds to time steps $i$ and $i+\Delta t$. To compute $\alpha$, we calculate the dot product of the unit vectors, $\alpha_i = \cos^{-1}(\vec{u}_i \cdot \vec{u}_{i+\Delta t})$ where $\vec{u}_i \cdot \vec{u}_{i+\Delta t} = \cos(\alpha_i)$.

Once the value of $\alpha$ is computed we can determine the angular displacement, $d = R \cdot \alpha_i$, where $R$ is the envelope radius. The squared displacement is $\Delta d^2$, that is the one we compute in the *gMSD* formula.

**Diffusion Coefficient Estimation:** From the gMSD curve, the translational diffusion coefficient $D_t$ is extracted using theoretical formulations:

$$\mathrm{gMSD}(t) = 4D_t t \quad \Rightarrow \quad D_t = \frac{\mathrm{gMSD}(t)}{4t}$$

### IX. DNA-PAINT SUPER-RESOLUTION IMAGING OF HIV PARTICLES

#### A. Cell Culture

All cells were maintained at 37°C in a humidified incubator with 5% $CO_2$. Culture media did not contain antibiotics. Lenti-X 293T producer cells (Takara Bio, 632180) cells were cultured in phenol-red-free DMEM/F12 medium (Thermo, 21041033) supplemented with 10% fetal bovine serum (FBS, Thermo).

#### B. Plasmid DNAs

Plasmids *pR8ΔEnv* (encoding the HIV-1 genome with an Env deletion) and *pcRev* (expressing HIV-1 Rev) were generously provided by Dr. Greg Melikyan (Emory University, Atlanta, GA,

USA). The *pCAGGS* plasmid encoding JR-FL Env was a kind gift from Dr. James Binley (Torrey Pines Institute for Molecular Studies, USA).

An ALFA-tag sequence (SRLEEELRRRLTE) was inserted into the variable loop 4 (V4) region of JR-FL Env, which lacked its C-terminal domain (Δ CT). The insertion was achieved using a two-step PCR reaction converging at the ALFA-tag site, as described in Williamson et al. (2025, *bioRxiv*).

#### C. Virus Production

Lenti-X 293T cells were seeded at a density of $6.5 \times 10^4$ cells/cm$^2$ in phenol-red-free DMEM/F12 supplemented with 300 nM Saquinavir mesylate (Sigma-Aldrich), an HIV-1 protease inhibitor, to produce immature virions. On the following day, the medium was replaced with FluoroBrite DMEM (Gibco, A1896701) containing 3% FBS.

Transfection was performed using GeneJuice (Merck, 70967) at a GeneJuice:DNA ratio of 3:1. The DNA mix consisted of *pR8ΔEnv*, *pcRev*, *Gag-FP-Delta Env* (where FP denotes a fluorescent protein, e.g., mTurquoise2), and *Env* in a molar ratio of 1:2:0.1:2.

For large-scale production, cells were plated in T-175 vented flasks and transfected in 24 mL of FluoroBrite medium using 12 µg of total DNA and 36 $\mu$L of GeneJuice diluted in 900 $\mu$L OptiMEM (Gibco, 11058-21). Culture supernatants were collected 48 hours post-transfection, centrifuged at 2000 × g for 5 minutes to remove debris, and filtered through a 0.45 $\mu$m syringe filter prior to concentration.

#### D. Virus Concentration

Virions were purified and concentrated by low-speed centrifugation through a sucrose cushion. After collection, the medium volume (typically 24 mL per T-175 flask) was adjusted to 28 mL with sucrose-free buffer (50 mM Tris-HCl, pH 7.4, 100 mM NaCl, 0.5 mM EDTA). This was underlaid with sucrose cushion buffer (50 mM Tris-HCl, pH 7.4, 100 mM NaCl, 0.5 mM EDTA, and 292 mM sucrose, i.e., 10% w/v) at a 4:1 (v/v) ratio.

Samples were centrifuged at 10,000 × g for 3–4 hours at 4°C with minimal acceleration and deceleration. Following centrifugation, the supernatant was carefully discarded, and tubes were inverted for 2 minutes to drain residual liquid. The resulting pellet, containing virions, was overlaid with 200–250 $\mu$L PBS (per T-175 pellet) and incubated for 30 minutes at 4°C.

Pellets were gently resuspended using pipette mixing, aliquoted into 20 $\mu$L portions, and stored at $-80$°C. For experimental use, virions were thawed on ice and directly used for sample preparation.

#### E. DNA-PAINT Sample Preparation

Glass-bottom 6-channel Ibidi $\mu$-slides (IB-80607) were first treated with 1 M potassium hydroxide (KOH) for 2 hours at room temperature to clean the surface, followed by thorough rinsing with ultrapure water. After air drying, the slides were exposed to ultraviolet (UV) light for 30 minutes to enhance surface activation. Viral samples were diluted 1:10 in phosphate-buffered saline (PBS) containing 2% paraformaldehyde (PFA) and incubated for 10 minutes at room temperature for fixation. A second 1:10 dilution in PBS was performed prior to applying the sample onto the glass surface, followed by a 30-minute incubation. Unbound material was removed by washing with PBS for approximately 5 minutes.

To minimize nonspecific binding, the surface was blocked for 30 minutes using PBS supplemented with 2% bovine serum albumin (BSA) and 0.2% fish skin gelatin. For labeling, a 25 nM solution of DNA-conjugated anti-ALFA-tag nanobody (MASSIVETAG-Q-ANTI-alfa; R1 sequence: 5′-TTTCCTCCTCCTCCTCCTCCT-3′ in 5% BSA/PBS was incubated on the surface for 1 hour at room temperature. Excess nanobody was removed by another 5-minute PBS wash, followed by a 5-minute incubation with 90 nm gold nanoparticles (G-90-100, CytoDiagnostics) for drift correction. A final PBS wash completed the sample preparation.

For imaging, the DNA imager strand (sequence R1: AGGAGGA) labeled with a 3′ Cy3b fluorophore was prepared in $C^+$ buffer (PBS with 1 mM EDTA, 500 mM NaCl, 0.02% Tween-20, pH 7.4). A final imager concentration of 1 nM was used for total internal reflection fluorescence (TIRF) microscopy.

#### F. TIRF Microscopy Setup

Imaging was performed on a custom-built TIRF microscope based on the Nikon Eclipse Ti-2 platform, equipped with a 100×/1.49 NA oil immersion objective and a Perfect Focus System. Excitation was provided by a 560 nm laser (1 W, MPB Communications), expanded and collimated using a variable beam expander and telescope. The beam was reshaped into a flat-top profile using a piShaper 6_6_VIS (AdlOptica).

Beam polarization was adjusted to circular using a linear polarizer and quarter-wave plate, then directed to the objective back focal plane via a lens and appropriate dichroic mirror (Di03-R405/488/561/635, Semrock). Emission light was filtered (FF01-446/523/600/677-25, Semrock) before being detected with a Hamamatsu ORCA-Fusion BT sCMOS camera. The resulting effective pixel size was 130 nm (after 2×2 binning).

#### G. DNA-PAINT Acquisition and Super-Resolution Analysis

Initial reference images were acquired using a 488 nm continuous-wave laser (100 ms exposure, 0.9 kW/cm$^2$) to localize Gag-mAmetrine viral markers. DNA-PAINT imaging was then performed, recording 40,000 frames at 100 ms per frame under 165 W/cm$^2$ illumination.

Super-resolution image reconstruction was carried out using the *Picasso* software suite. Fiducial markers (gold nanoparticles) enabled lateral drift correction via redundant cross-correlation, with approximately 50 fiducials per field of view ensuring accurate alignment. Final images were rendered as Gaussian localizations, color-mapped by localization precision. Co-registration with diffraction-limited Gag signals confirmed viral specificity.

#### H. Quantitative PAINT (qPAINT) and Cluster Analysis

Fluorescent time series for each cluster were analyzed using custom MATLAB (v2022a) scripts implementing the qPAINT method to quantify protein copy numbers per cluster. Clusters were identified using the DBSCAN algorithm with an epsilon ($\varepsilon$) of 5 nm and a minimum of 10 localizations (*minPts*), optimized to resolve Env protein clusters.

For each cluster, the temporal gaps (dark times) between localizations were extracted and used to generate cumulative histograms. These were fitted with exponential decay functions to obtain the mean dark time $\tau_0$, from which the qPAINT index $\xi = \tau_0^{-1}$ was calculated.

Calibration was performed by analyzing small clusters (maximum pairwise point distance $\leq$ 55 nm) to determine the baseline $\xi$ corresponding to a single binding site, identified as 0.0096 s$^{-1}$. The number of Env proteins per cluster was estimated as the ratio $\xi/\xi_{\text{single}}$. Protein locations were assigned to individual localizations using K-means clustering, with the number of clusters $K$

determined from qPAINT-based protein quantification.

#### I. Experimental and simulations comparison results

To complete the cluster analysis, we use the data observed in FIG 3 and FIG 4 for experimental. In FIG 5 we have the plots for our simulations with 12 and 24 spike proteins.

### X. TRANSLATIONAL DIFFUSION OF SPIKE PROTEINS OVER THE ENVELOPE

We compute the $D_{sp}$ within the range $1–10, \tau_v$. Across all spike counts, the gMSD curves exhibit similar behavior, initially linear $\alpha = 1$, before transitioning to a plateau, indicating a saturation regime. The values of $D_{sp}$ remain relatively consistent across all three configurations, as expected in confined diffusion. Given the diffusion coefficient $D_{sp}$, we define the saturation time as: $\tau_{\text{sat}} = R^2/D_{sp} = 90\tau_{\text{sdpd}}$.

As a theoretical case, we can compute the Stokes-Einstein translational diffusion over a surface,

$$D_t = \frac{k_B T}{4\pi\eta r_s} = 0.039.$$

TABLE IV. Spike translational diffusion coefficients.

| **Number of spikes proteins** | $D_{sp}[l^2_{\text{sdpd}}/\tau_{\text{sdpd}}]$ | $\tau_{sat}[\tau_{\text{sdpd}}]$ |
|---|---|---|
| Stokes-Einstein 2D | 0.039 | 100 |
| 2 | $0.044 \pm 2\times10^{-3}$ | 90.5 |
| 12 | $0.045 \pm 2\times10^{-4}$ | 89.3 |
| 24 | $0.045 \pm 3\times10^{-5}$ | 89.5 |

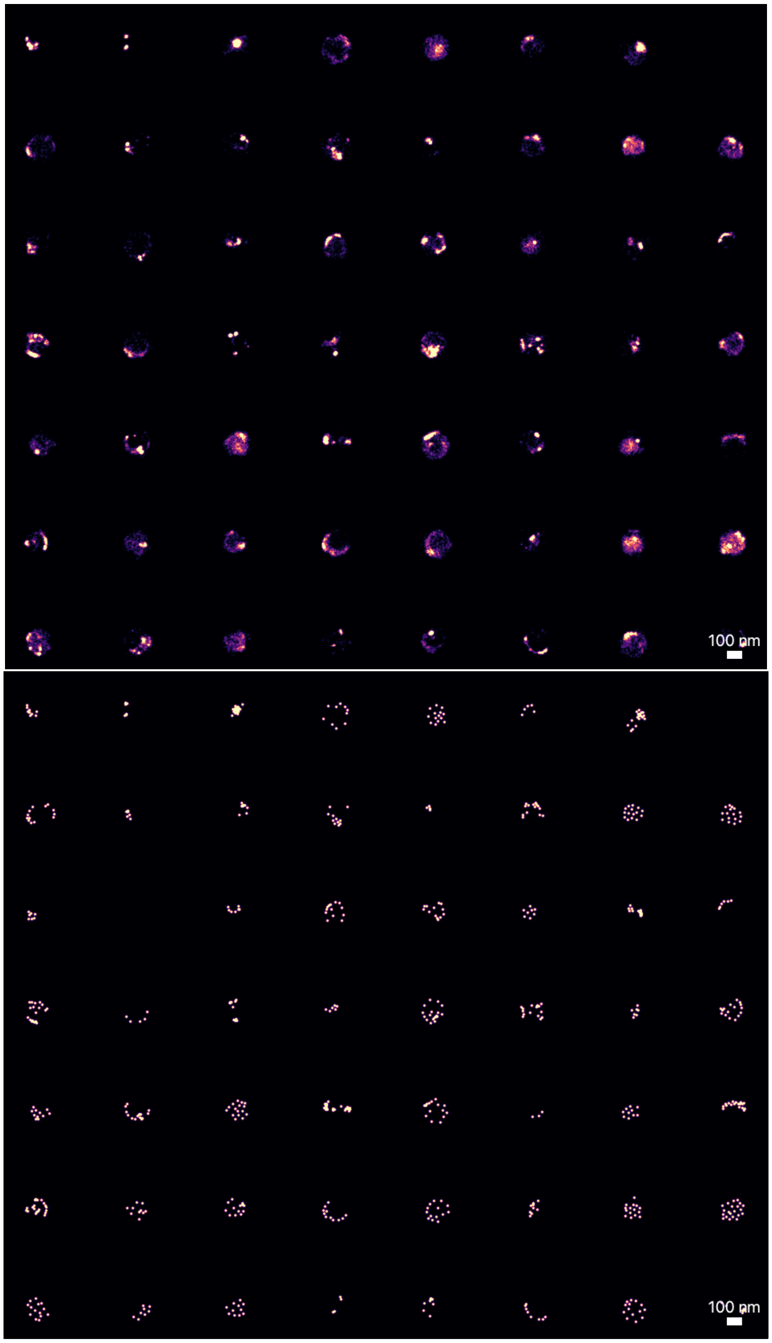


FIG. 3. HIV-1 immature viruses decorated with JR-FL-v4-ALFA-tag (DeltaCT) / Gag-mTurquoise2. In a) DNA-PAINT. In b) Proteins projections.

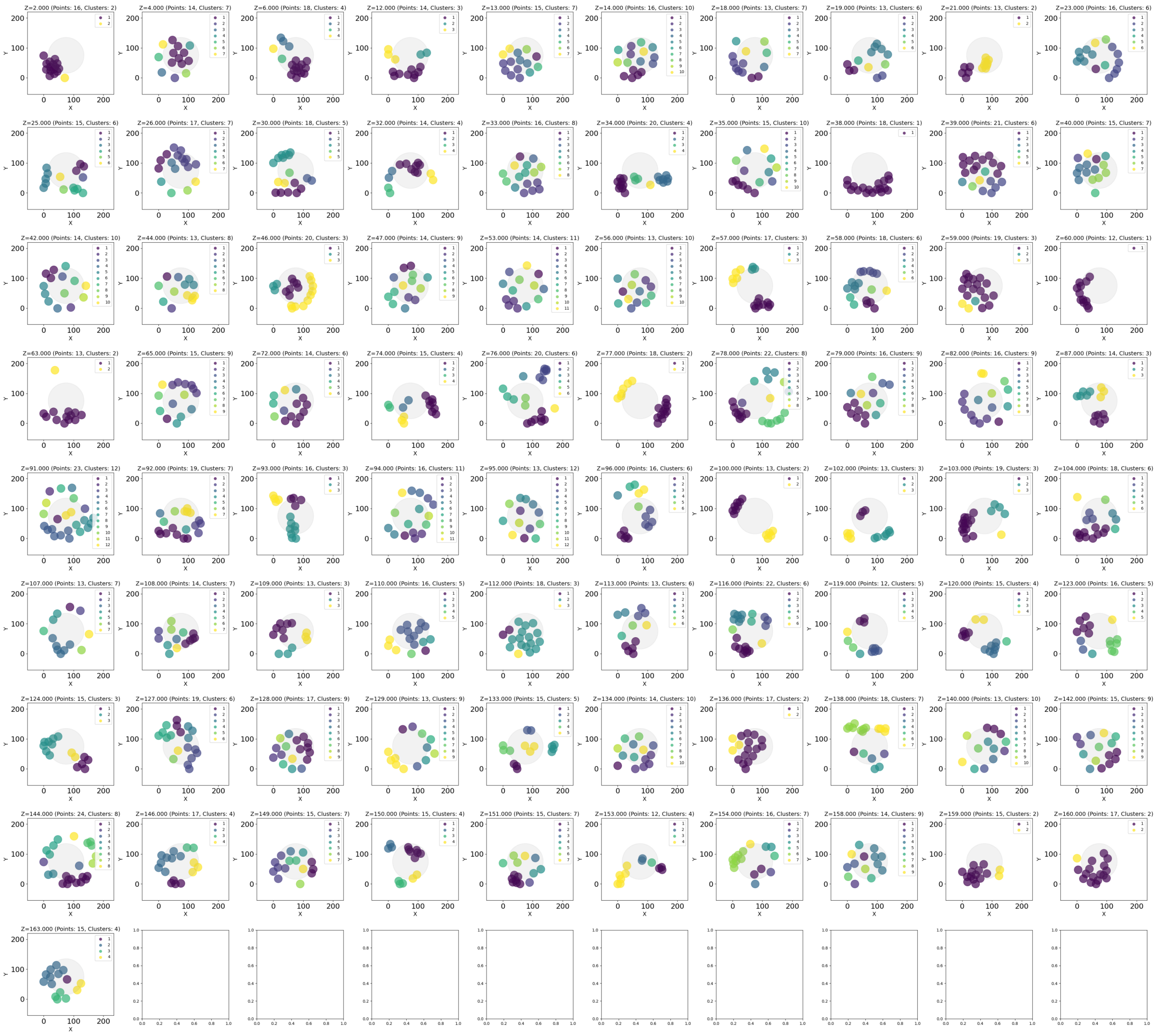

FIG. 4. Experimental HIV-1 immature viruses filtered from 12 to 24 spike proteins

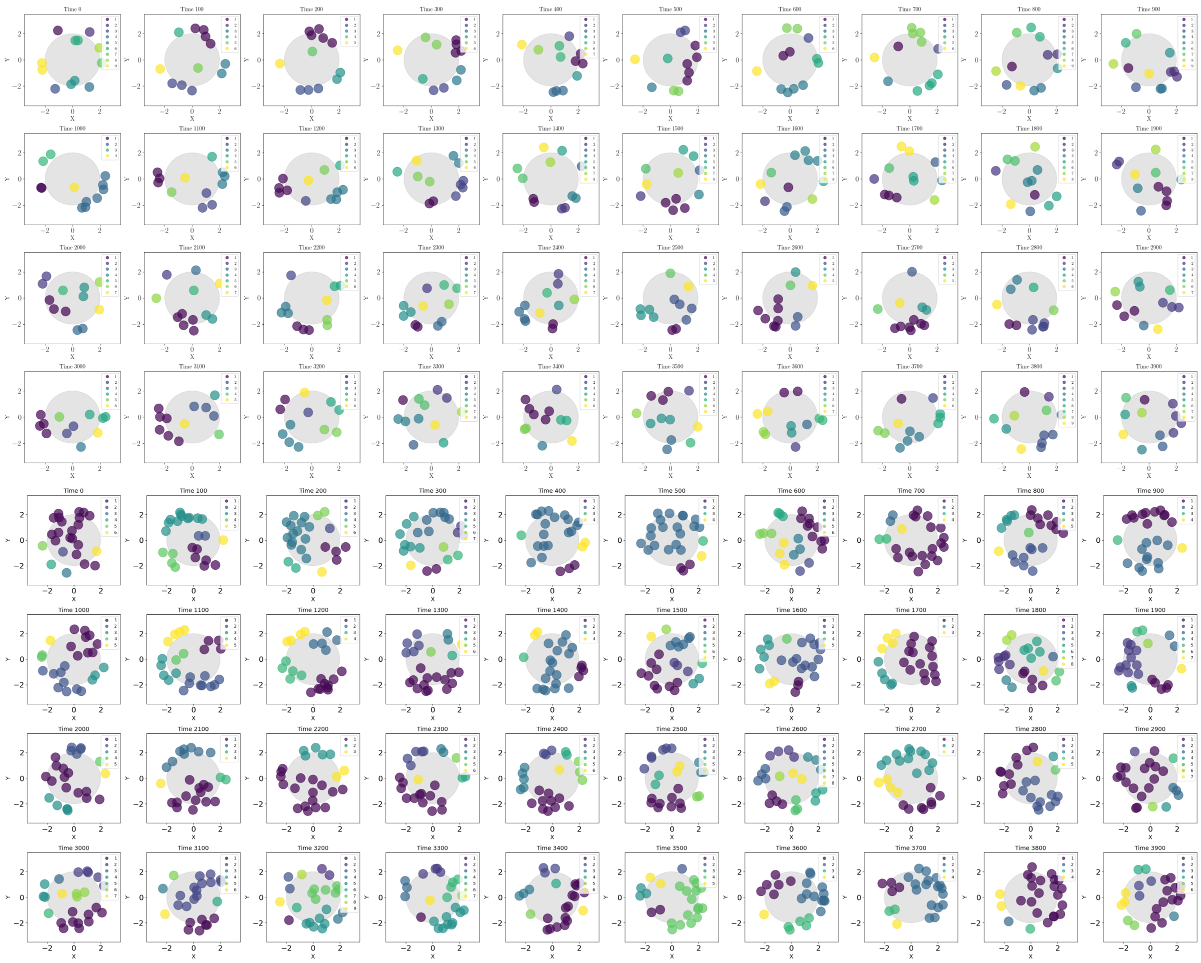

FIG. 5. Simulation - 12 and 24 spike proteins

[1] J. Padding and A. Louis, Physical Review E-Statistical, Nonlinear, and Soft Matter Physics **74**, 031402 (2006).

[2] J. J. Monaghan, In: Annual review of astronomy and astrophysics **30**, 543 (1992).

[3] A. Ledesma-Durán and L. H. Juárez-Valencia, European Physical Journal E **46** (2023).

[4] A. Vázquez-Quesada and M. Ellero, SeMA Journal **79**, 165 (2022).

[5] M. Linke, J. Köfinger, and G. Hummer, The Journal of Physical Chemistry Letters **9**, 2874 (2018).

[6] A. Ledesma-Durán, J. Munguía-Valadez, J. A. Moreno-Razo, S. I. Hernández, and I. Santamaría-Holek, Frontiers in Physics **9** (2021).
